# Evolutionary determinants of reproductive seasonality: a theoretical approach

**DOI:** 10.1101/2022.08.22.504761

**Authors:** Lugdiwine Burtschell, Jules Dezeure, Elise Huchard, Bernard Godelle

## Abstract

Reproductive seasonality is a major adaptation to seasonal cycles and varies substantially among organisms. This variation, which was long thought to reflect a simple latitudinal gradient, remains poorly understood for many species, in part due to a lacunary theoretical framework. Because seasonal cycles are increasingly disrupted by climate change, a better understanding of the ecology of reproductive seasonality could generate important insights on how climate change may affect biodiversity. The goal of this study was to investigate the drivers of evolutionary transitions towards reproductive seasonality using a realistic agent-based optimisation model simulating the life cycle of a female yellow baboon, who typically breeds year-round. Specifically, we tested the influence of three ecological traits (environmental seasonality, productivity and unpredictability) and three life-history traits (daily reproductive energy expenditure, reproductive cycle length and infant extrinsic mortality) on the intensity of reproductive seasonality. To do so, we simulated diverse reproductive phenology strategies (from non-seasonal to highly seasonal), assessed which were optimal and computed, for the set of optimal strategies, the intensity of reproductive seasonality. We then induced variation in each trait of interest and examined how it affected the intensity of reproductive seasonality. We found significant effects of all three environmental traits: high reproductive seasonality was favoured by high environmental seasonality, low environmental productivity and low unpredictability. It was further, and most strongly, favoured by high daily reproductive energy expenditure. In contrast, there was no significant effect of reproductive cycle length and infant extrinsic mortality. Our modelling approach successfully disentangled the effects of environmental seasonality, productivity and unpredictability on the intensity of reproductive seasonality, which likely all contribute to generate the well-known association between latitude and reproductive seasonality. Our results further highlight the critical importance of life history pace on the evolution of reproductive seasonality. Overall, this study contributes a powerful theoretical framework and modelling tool that may apply across the life-history space, as well as sheds new light on the emergence and maintenance of non-seasonal breeding in slow-living species, including humans.

## Introduction

Reproductive seasonality, which consists in a temporal gathering of reproductive events, is a major adaptation to seasonal cycles as it allows synchronising the energetic costs of reproduction with the annual food peak (van Schaik & Brockman, 2005). Because seasonal cycles are increasingly disrupted by climate change (Easterling *et al*., 2000), a better understanding of the ecology and evolution of reproductive seasonality could generate important insights on how climate change may affect biodiversity (Thackeray *et al*., 2016).

Many studies have focused on understanding the physiological regulation of reproductive seasonality as well as variation in its timing (Brockman & van Schaik, 2005; Bradshaw & Holzapfel, 2007; Bronson, 2009; Vatka *et al*., 2014; Clauss *et al*., 2021), but less work has investigated why the intensity of reproductive seasonality exhibits such a broad range of variation across species (Campos *et al*., 2017). Understanding the drivers of such variation in strategies of reproductive phenology (i.e. timing and length of the annual birth season) is essential to predict the resilience of species to climate change.

The intensity of environmental seasonality (often approximated by latitude) has traditionally been used to predict reproductive seasonality (Conover, 1992; Ogutu *et al*., 2014; Heldstab, 2021b) with tropical species breeding year-round and temperate and Arctic species breeding seasonally. Yet, a great part of its variation remains unexplained by this predictor alone. For instance, some tropical species reproduce seasonally (Brown & Shine, 2006; Hongo *et al*., 2016) while non-seasonal breeding has been observed in some temperate (Riedman *et al*., 1994) and Arctic (Gruyer *et al*., 2010) species. There are also examples of sympatric species which largely differ in their reproductive phenology (Sinclair *et al*., 2000). In sum, besides environmental seasonality, other determinants likely play a significant role in the evolution of reproductive seasonality, such as other ecological factors, life-history traits, and their interaction (English *et al*., 2012; Clauss *et al*., 2021)

Ecological factors other than environmental seasonality could include environmental productivity (average level of food availability) and unpredictability (hereafter approximated by the non-seasonal variation in food availability and modelled as an added noise to food availability). First, in highly productive habitats, the energy requirements of reproduction could be met year-round even in seasonal environments, thus decreasing the benefits of seasonal breeding. Interestingly, environmental seasonality and productivity are both highly correlated with latitude (Botero *et al*., 2014), which is a major predictor of reproductive seasonality (Heldstab, 2021b; Heldstab *et al*., 2021). Yet, most studies only investigate the global effect of latitude without disentangling the effects of environmental productivity, seasonality and their interaction. Second, in unpredictable environments, even with some degree of environmental seasonality, a flexible reproduction may be more advantageous than a strictly seasonal reproduction. Indeed, reproducing seasonally in an environment where the food peak is regularly reduced or delayed may generate mismatches between a fixed reproductive timing and a variable annual food peak (Vatka *et al*., 2014; Clauss *et al*., 2021). Such mismatches could reduce the benefit of breeding seasonally and therefore lead to a decrease in the intensity of reproductive seasonality. However, only a few studies have investigated this latter effect of environmental unpredictability on the *intensity* (and not only on the *timing*) of reproductive seasonality, with mixed results so far. While English *et al*. (2012) found that inter-annual variation in food availability had an effect on the intensity of birth synchrony of wild ungulate populations from 38 species, two other studies on red deer, *Cervus elaphus* L. (Loe *et al*., 2005) and chacma baboons, *Papio ursinus* (Dezeure *et al*., 2023) found no effect of environmental unpredictability on reproductive seasonality.

Regarding the effect of life-history traits, variation across species in strategies used to finance the energetic costs of reproduction has likely played a major role in shaping the evolution of their reproductive schedule. In particular, the mechanism consisting in limiting the peak in energy demand for reproduction, by adopting a “slower” life history, as defined by the fast-slow continuum framework (Stearns, 1989; Bielby *et al*., 2007) could be critical to sustain non-seasonal reproduction. Indeed, high day-to-day reproductive energy requirements, which are typical of short-lived species with large litter size and fast reproductive cycles (Bielby *et al*., 2007), likely limit the period of the year where the energetic costs of reproduction can be met. In contrast, in species with slower reproductive paces, reproductive costs are spread over time and more easily afforded year-round (van Noordwijk *et al*., 2013). Additionally, and regardless of the life-history pace, the length of the reproductive cycle could also be a key factor, depending on whether or not it is synchronised with the annual schedule. Particularly, for long lived species, a reproductive cycle lasting more than exactly one, two, three or more full years (i.e., that is not an integer number of years), could decrease the benefits of seasonal breeding by introducing gaps at the end of each reproductive cycle, where females must wait the next reproductive season in order to reproduce again. Lastly, slow-living species often have small litters or singletons that can be lost all at once (Bielby *et al*., 2007). In such cases, species with high external offspring mortality compared to that of adults – a defining feature of slow life histories (Jones, 2011) – could face costs from reproducing seasonally by experiencing gaps after losing their offspring if they have to wait the next reproductive season to conceive again, similarly to species whose reproductive cycles are not integer years.

Finally, despite an abundant empirical literature on reproductive seasonality, few models have been developed to explain the variation of reproductive strategies in seasonal environments (Tökölyi *et al*., 2012; Stephens *et al*., 2014; Kristensen *et al*., 2015), with only one, to our knowledge, studying the emergence of reproductive seasonality (Sun *et al*., 2020). Yet, this last model focuses on the evolution of income versus capital breeding and considers environmental seasonality as the only determinant of the emergence of seasonal breeding. This highlights a general lack of theory regarding the drivers of the intensity of reproductive seasonality, and a specific need to develop a predictive framework integrating the combined effects of various ecological and life-history traits across ecological and evolutionary timescales. In addition, because several potential determinants of the intensity of reproductive seasonality typically co-vary in natural environments, modelling approaches appear essential to disentangle their effects by introducing variation in some traits while keeping others constant.

In this study, we developed an agent-based optimisation model to investigate the effects of variations in ecological and life history traits on the emergence of reproductive seasonality. We chose to create a realistic and detailed model based on energetics, that we parameterized using published empirical data from the yellow baboon (*Papio cynocephalus*). This choice of a long-lived tropical and non-seasonally breeding (Campos *et al*., 2017) model species contrasts with most studies on reproductive seasonality. First, focusing on a non-seasonal breeder and on how reproductive seasonality could emerge from this point is a new approach and the *Papio* genus is well suited for it because it is characterised by phenological flexibility, with one in six *Papio* species breeding seasonally (Petersdorf *et al*., 2019). Second, studying a tropical species like baboons is of major interest because most studies of reproductive seasonality focus on organisms from temperate regions (Bronson, 1985; Vatka *et al*., 2014). Lastly, long-lived species such as baboons, where reproductive cycles spread over multiple years, tend to be under-represented in studies of breeding seasonality, but could bring important insights to understand our own reproductive phenology, namely why humans reproduce year-round.

Yellow baboons live in tropical semi-arid savannas characterised by relatively low productivity, high unpredictability and moderate seasonality (Gesquiere *et al*., 2008). They give birth to one offspring at a time with a limited growth rate during lactation (Altmann & Alberts, 1987) inducing relatively low day-to-day reproductive energy requirements. Their mean interbirth interval (IBI) is 638 days, or 1.75 years (Gesquiere *et al*., 2018), which is not an integer number of years, and they exhibit a moderate extrinsic infant mortality (for a slow-living species) (Alberts, 2019). Altogether, their non-seasonal reproductive phenology appears consistent with our hypotheses regarding their habitat characteristics and life history traits.

We therefore simulated variable strategies of reproductive phenology, from non-seasonal breeding to a highly seasonal reproduction, together with their energy budget, in order to calculate their associated fitness outcome. The phenology strategies that maximized individual fitness, here calculated as λ_ind_, the individual-level equivalent of the population growth rate (McGraw & Caswell, 1996), were interpreted as the optimal reproductive phenology strategies. We first identified the optimal reproductive phenology strategies in the natural environment of yellow baboons, the Amboseli National Park, source of most empirical data used here. To do so, we used the Normalized Difference Vegetation Index (NDVI), a satellite-derived quantitative measure of vegetation productivity, extracted from the Amboseli national Park coordinates, as a proxy of food availability. By artificially reshaping the NDVI time series, we subsequently investigated the effect of variation in three environmental traits (seasonality, productivity and unpredictability) on the fitness of these different strategies. We further tested the influence of three life-history traits (daily reproductive expenditure, length of the reproductive cycle and infant extrinsic mortality rate) on the optimal phenology strategy by modifying the model parameters. By doing so, we were able to test our working hypotheses, respectively proposing that the emergence of reproductive seasonality is favoured by (H1) high environmental seasonality, (H2) low environmental productivity, (H3) low environmental unpredictability, (H4) high daily energetic expenditure of reproduction, (H5) a reproductive cycle that is an integer number of years and (H6) low infant extrinsic mortality rate.

## Methods

### A. Model description

The programme code uses a combination of R (R Core Team, 2020) and C++ functions through the Rcpp package (Eddelbuettel & François, 2011). The model description follows the ODD (Overview, Design concepts, Details) protocol for describing individual and agent-based models (Grimm *et al*., 2010).

#### 1. Purpose

The purpose of this study is to investigate the determinants of reproductive seasonality in a mammalian species. To do so, we simulate the life cycle of a female yellow baboon (*Papio cynocephalus*) with an energetic approach to evaluate the optimal strategy of reproductive phenology. We parameterized the model using published empirical data (Table S1), but we developed its structure to be as general as possible so that it could be adapted to other mammalian species.

#### 2. Entities, state variables, and scales

##### Agents/individuals

The model simulates the life cycle of a female adult baboon and her offspring. We do not discriminate the offspring by sex and only include adult females. We consider here that males are non-limiting for female reproductive success, and do not affect the outcome of reproduction, which sounds reasonable in a species where paternal care is optional (Buchan *et al*., 2003). Therefore, each individual in the model belongs to one of four possible life stages: foetus, infant, juvenile and (adult) female. A foetus is created when a female conceives and switches to the next life stages through defined events: birth, weaning and sexual maturity. As individuals grow, they acquire new abilities (e.g. foraging or reproducing). Technically, we simulated these individuals using object-oriented programming where the four life stages are represented by four classes that inherit (acquire properties and behaviour) from one another (Fig. 1). Each individual is characterised by some attributes or state variables, according to his life stage (Appendix A, Table S2) which accounts for the changes occurring during the simulation.

**Figure 1:**
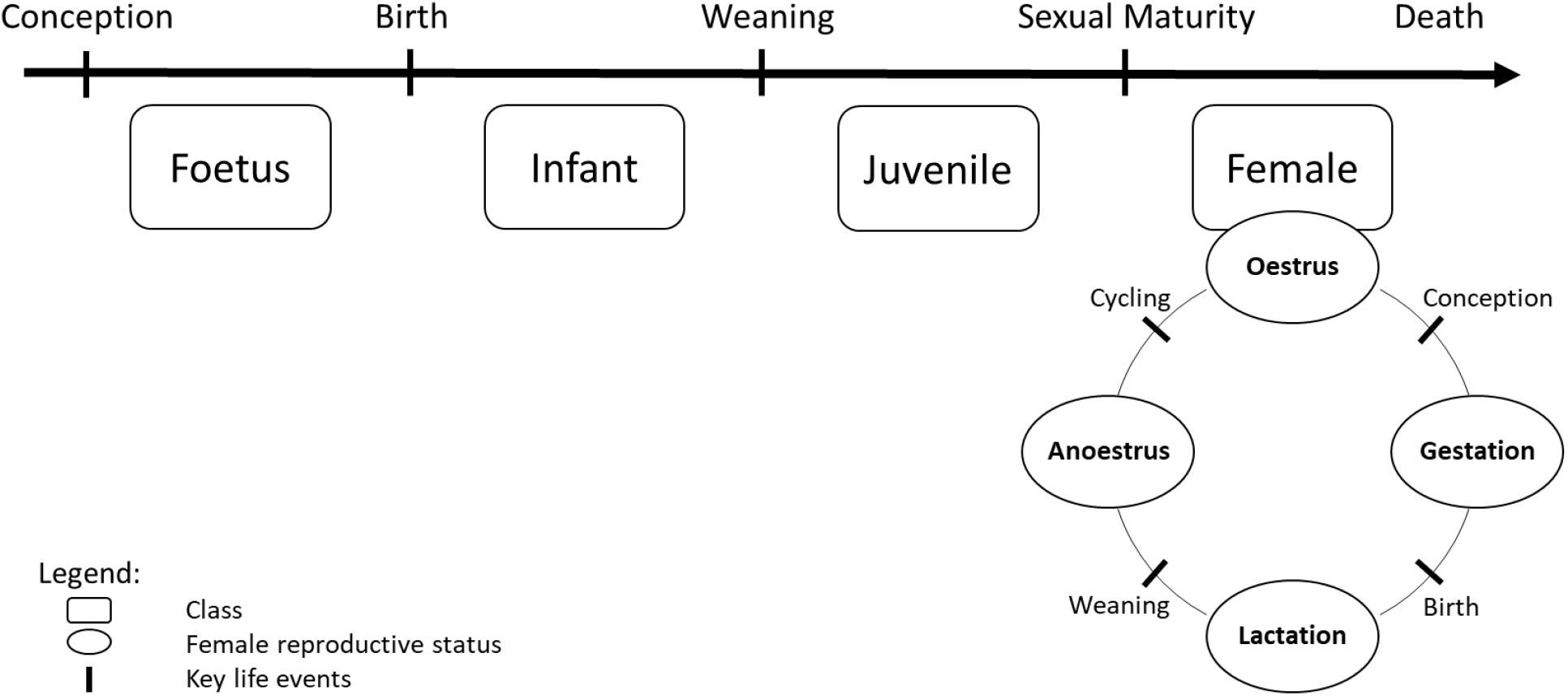
Schematic representation of the class-structure used in the model. Each rounded rectangle represents a class. The Foetus class is the base-class while the Female class is the class with the most advanced properties, including reproduction. Class Female individuals can therefore produce offspring that will appear as an individual of class Foetus and evolve to the other classes through key life events associated with changes in the female reproductive status.

##### Temporal scales

One time step represents one day and simulations end with the female death or her last offspring reaching sexual maturity, depending on which event occurs last.

##### Environment

Each simulation occurs in an environment characterised by its food availability, approximated by the Normalized Difference Vegetation Index (NDVI). NDVI is an indicator of vegetation productivity (Pettorelli *et al*., 2005) and is therefore relevant to predict food availability for yellow baboons whose diet, although omnivorous, consists mainly in plants (Post, 1982). NDVI has indeed previously been used in several baboon populations to estimate food availability (Baniel *et al*., 2018; Dezeure *et al*., 2021; Walton *et al*., 2021; Dezeure *et al*., 2023) and has also proved relevant to predict reproductive seasonality in humans (Macfarlan *et al*., 2021). When vegetation productivity is weak (low NDVI), the model also allows the use of alternative sources of food, called ‘fallback food’, as described in wild yellow baboons (Altmann, 1998) (see Appendix B - Section 3 “Energy intake” and Figure S1 in the Supplementary Material for details).

To account for realistic seasonal changes in NDVI, we used real values from the Amboseli population of yellow baboons. We used a GPS tracking study (Amboseli Baboon Research Project, 2015 - see Markham *et al*., 2013 for methods) to assess the coordinates of a quadrat where the majority of the baboon movements occurred (South/West: -2.75, 37.04 ; North/East: -2.70, 37.11). This quadrat covers a surface of about 42 km², which is consistent with the annual home range size observed in wild baboons (Altmann & Muruthi, 1988). We extracted NDVI within this quadrat using the 250-m resolution NDVI 16-day composite data from the MOD13Q1 product (Didan, 2015) which we linearly interpolated to have a daily time series from February 18, 2000 to May 09, 2022. Because our simulations can run for more than 27 years, we needed to extend the NDVI dataset. To do so, we chose to simply repeat the entire NDVI time series. To minimise discontinuity, we compared the first and last years of the time series and selected the day of year with the least variable NDVI that we therefore used as the transition point for concatenation of the NDVI time series.

#### 3. Process overview and scheduling

The same sequence happens at every time step (day) of the simulation.

1. Potential death of the female and/or offspring (if any)
2. Potential change of the female reproductive status (cycling, conception, birth or weaning)
3. Evaluation of daily energy intake extracted from the environment
4. Evaluation of daily energy needs based on reproductive status
5. Energy allocation:

5.1. If positive energy balance: storage of energy for the female and offspring (if any)
5.2. If negative energy balance: release of energy and/or energy restrictions (slowdown of growth, miscarriage, infant abandon) for the female and offspring (if any)
6. Growth of the female and/or dependent offspring (if any)

Following the death of a female, the same sequence (without reproduction) is repeated for each independent offspring (juvenile) until they die or reach sexual maturity. Dependent offspring (foetus or infant) do not survive after their mother’s death.

These six submodels (Death, Change of reproductive status, Energy intake, Energy needs, Energy allocation and Growth) are described in Appendix B of the supplementary material (Fig S1-3).

#### 4. Design concepts

##### Emergence, Adaptation

In this statistic optimisation model, we compute the female fitness associated with each phenology strategy. The fitness of an individual does not depend on the strategy of others. In particular, male function is supposed to be non-limiting, and we do not consider density-dependence interactions. The phenology strategy (i.e. the beginning and length of the time window during which a female can conceive) is constant throughout each simulation (i.e. throughout her lifetime), but different phenology strategies are tested in a given set of simulations. To account for both different reproductive timings (the date in the annual cycle of the mating season) and different intensities of the reproductive seasonality (i.e. the length of the mating season), we tested various phenology strategies, with reproductive windows starting each first day of each month (12 different timings) and lasting from 1 to 11 months (11 different intensities of reproductive seasonality). We also tested the non-seasonal phenology strategy (females can conceive year-round), resulting in 133 phenology strategies tested for each set of simulation. We ran 2000 simulations for each phenology strategy and extracted the resulting mean fitness values. The optimal strategies of reproductive seasonality emerged from the set of simulations as the strategies associated with the highest fitness values. Because differences between some phenology strategies can be small (one-month difference in start or length of reproductive window), in most cases there is not a single phenology strategy that emerges but rather a group of optimal phenology strategies associated with the highest fitness values. More precisely, we assessed the difference between the mean fitness value of the “best” phenology strategy (i.e. highest one) and the mean fitness values associated with each other phenology strategy and allowed a 5% decrease in fitness as a confidence margin. Optimal phenology strategies were therefore defined as strategies whose mean fitness value ranged above 95% of the highest mean fitness value.

##### Fitness

Individual fitness (λ_ind_) is calculated at the end of each simulation as described in McGraw and Caswell (1996) and represents the “population growth rate of an individual”. Previously used in empirical studies (McLean *et al*., 2019), this value takes into account not only the number of offspring born to a female but also the timing at which they are produced: the earlier in life a female has offspring, the higher her individual growth rate. Specifically, λ_ind_ is the dominant eigenvalue of Leslie matrices built on individual females, which indicates the number of offspring produced each year. It should be noted that λ_ind_ does not represent an individual’s contribution to the demographic growth of the population. In particular, the mean of several individual fitness values cannot be used to estimate a population growth rate (McGraw & Caswell, 1996). Because our simulations begin when the female has just reached sexual maturity, we decided to consider in the calculation of the individual fitness only offspring having reached the same point (i.e. sexual maturity). This method prevents us from overestimating or underestimating any life stage in the calculation of fitness, and it especially allows to take into account the post-weaning period which is known to be critical for juveniles (McLean *et al*., 2019). Even if sexual maturity seems to appear around 10 months later for male offspring than for female offspring in yellow baboons (Charpentier *et al*., 2008), early maturation has been observed in male hamadryas baboons (Zinner *et al*., 2006). Because of the uncertainty on male sexual maturity age and for simplicity, we chose to use the female age at sexual maturity as a unique milestone for offspring of both sexes.

##### Stochasticity

There are two stochastic processes in this model: conception and extrinsic mortality. Variation in the probability of conception is accounted for by randomly picking a cycling duration from observed data in the Amboseli yellow baboons (Gesquiere *et al*., 2018) before each cycling phase of the reproductive cycle. Extrinsic mortality (external causes of death) is accounted for by assigning a maximal lifespan to each individual, also randomly picked from empirical data (McLean *et al*., 2019) and independent of environmental variations. Individuals die if they reach their maximal lifespan, independently of their current energy resources. Yet, individuals can also die before their assigned lifespan is reached, if the environmental conditions do not provide them with enough energy to survive (“intrinsic” mortality). Overall, individuals in the model can either die from a stochastic external mortality or from an environmentally-dependent intrinsic mortality (see the submodels “Death” and “Cycling” in supplementary for details). Finally, the first day of the simulation is also randomly picked in the annual cycle, in order not to favour one time of the year over another. We ran each simulation 2000 times to account for such stochasticity. We assessed this number by increasing the number of simulation runs until reaching stability in the results.

##### Observation

The model outputs recorded at the end of a simulation are:

1. The fitness of the female with respect to the phenology strategy she has followed
2. The seasonality of births that took place during the simulation

We used circular statistics to characterise the seasonality of births, according to recent recommendations (Thel *et al*., 2022). In circular statistics, days of the year are represented as angles on a circle and each birth event is characterised by a vector of length 1 pointing to the day of year when it occurred. By computing the mean vector of all birth events, we can characterise their seasonality: the direction (µ) of the mean vector gives the mean date of birth, while its length (r) defines the intensity of seasonality (r = 0 when births are evenly distributed and r = 1 when they all occur on the same day).

#### 5. Initialisation

At time t=0, only one individual of class Female is created. It represents a female yellow baboon that has just reached sexual maturity and its state variables are initialised using parameters from the literature (Appendix A, Table S3). Initial attributes for offspring are presented in Appendix A, Table S4.

#### 6. Input data

We use some species-specific parameters in the model that remain constant throughout each simulation. We compiled and calculated them from the literature (Table S1). Most of them come from published empirical data from wild populations of yellow baboons. When such data were unavailable, we used available values from phylogenetically close species or captive populations.

##### B. Test of hypotheses

To test our hypotheses (H1-6), we artificially modified in the model the value of each of the six environmental or life history traits investigated and computed the associated birth seasonality.

#### 1. Computation of simulated birth seasonality

To compute the birth seasonality associated with specific environmental and life history conditions, we first evaluated all 133 possible phenology strategies (from highly seasonal to non-seasonal) to identify which were optimal (i.e. associated with the highest fitness values, see the section “*Emergence, Adaptation*” section above). We then pooled all births coming from the pool of optimal phenology strategies, and calculated, from these births dates, the mean vector length (r) and direction (µ), using the functions ‘rho.circular’ and ‘mean.circular’ from the ‘circular’ package (Lund *et al*., 2017) as a measure of birth seasonality.

#### 2. Artificial variation of environmental and life history effects

In order to disentangle the effects of environment and life history, we chose to modify only the three environmental traits first. By doing so, we drew an environmental landscape composed of three axes (productivity, seasonality and unpredictability) in order to explore their whole range of (co)-variation. We expected one or more interactions, and specifically that the effect of environmental seasonality on reproductive seasonality would be modulated across a gradient of environmental productivity, with higher productivity extending the annual time window where reproduction is sustainable in seasonal environments. For example, abnormally extended birth seasons were observed in a population of North American elk that punctually experienced a higher habitat quality (Keller *et al*., 2015). Because a 3D representation of the environmental parameter space (productivity, seasonality and unpredictability) can be difficult to apprehend and increases computing time, we chose to systematically simulate and represent only a two-dimensional environmental landscape covering the full range of variation of both environmental seasonality and productivity – and later represented through heatmaps. Variation in those two parameters can indeed reflect a variation in latitude with the top-right to bottom-left diagonal of the heatmaps representing a gradient from low latitudes (high productivity and low seasonality) to high latitude (low productivity and high seasonality). We then tested for the effect of unpredictability, the third environmental dimension, by simulating this same heatmap for four levels of unpredictability (i.e. sectional views of the 3D environmental landscape). We subsequently tested the effect of each life history trait on reproductive seasonality across this two-dimensional landscape in order to cover a broad range of viable environments, but kept unpredictability unchanged for simplicity.

#### 3. Modification of environmental traits: test of hypotheses H1-3

In order to test the influence of environmental seasonality, productivity and unpredictability (H1-3), we first decomposed the NDVI time series into three components, therefore disentangling seasonal from non-seasonal variation (Fig. S4):

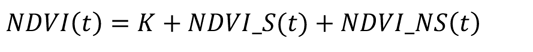

K is the mean NDVI over the entire time series and is therefore a constant that represents the overall productivity of the environment. NDVI_S represents the seasonal part of the variation in NDVI and is obtained by calculating, for each day of the calendar year, the mean daily offset of NDVI from its mean over the 22 years of data (i.e. the mean daily NDVI, minus K). NDVI_S captures the within-year variation of NDVI and is completely identical between years. Lastly, NDVI_NS captures the rest of the NDVI variation and is computed by removing to the overall NDVI its mean K and its seasonal component NDVI_S. NDVI_NS can be seen as the noise, where a positive value indicates that the NDVI is higher than expected for a given calendar day, i.e. for the season. In other words, NDVI_NS essentially captures the non-seasonal variation in NDVI, later used as a proxy for environmental unpredictability. It should be noted that NDVI_NS also encompasses the noise due to measurement errors, that cannot be disentangled from the biological noise but that is likely negligible compared to non-seasonal environmental variation. In addition, unpredictability is a multidimensional entity that cannot be restricted to the NDVI_NS component alone: for example, one additional dimension is the unpredictability in the timing of the food peak. In other words, this decomposition describes the changes in NDVI magnitude for the same calendar date, but does not directly capture the changes in timing of the NDVI time series.

We then artificially modified these different components of the NDVI time series by introducing three parameters, *s*, *p* and *u*. This allowed us to modulate the contribution of each component of NDVI as follow:

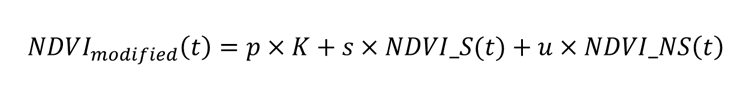

The higher *p*, *s* and *u*, the higher the environmental productivity, seasonality and unpredictability (respectively). The parameter *p* varied from 0.85 to 1.4 while *s* and *u* varied from 0 to 3, with the real NDVI given by *p* = *s* = *u* = 1. NDVI being a normalised index whose values range between 0 and 1, we truncated accordingly the modified NDVI time series, replacing any negative values with zero, and any value superior to one with one. The ranges of variation for the three parameters *p*, *s* and *u* were chosen in order to explore substantial variation in environmental productivity, seasonality and unpredictability while still simulating viable environments for baboons (i.e., with a sufficiently high productivity). Non-viable environments are characterised by a fitness of zero and appear as grey cells on heatmaps.

#### 4. Modification of life history traits: test of hypotheses H4-6

To investigate the effect of an increase in daily reproductive energy expenditure (H4), we modified the daily growth rate at 7.5 g/day instead of the original 5g/day, which is a highly variable parameter between species with different life-history paces (Bielby *et al*., 2007). This represents an increase of 50% in the energy expenditure associated with growth. Gestation was accordingly reduced from 178 days to 118 to ensure a constant birth mass.

To test the effect of a reproductive cycle being an integer number of years (H5), we chose to artificially modify the year length, instead of the reproductive cycle length. With this approach, the biological timing (and especially the daily reproductive energy expenditure) remains unchanged, but the temporal synchronisation between reproductive cycle and year length can change, thus disentangling the two effects predicted by H4 and H5. Reducing or extending the reproductive cycle length (for example by modifying gestation length, growth rate, weaning mass…) would indeed be an alternative approach to synchronise the reproductive schedule with the annual cycle. Yet, by also modifying the energetical aspects of the reproductive cycle, this approach could influence reproductive seasonality indirectly, for example via an effect of daily reproductive energy expenditure (H4). On the contrary, modifying the year length alone allows us to isolate the effect of synchronising the reproductive cycle with the annual cycle without altering the energetic transfers between mother and offspring. To do so, we interpolated the 16-days NDVI time series from the raw data assuming a year length of 637 days (equal to the interbirth interval), 425 days (so the IBI would be of exactly 1.5 year) and 365 days (normal conditions, IBI = 1.75 year). In other words, we “stretched” the NDVI time series so its seasonal variations would be perfectly synchronised with the reproductive cycle (year length = interbirth interval) or desynchronised (year length = 1.5 interbirth interval). For example, in the synchronised configuration, with a year length of 637 days, the annual good season would always fall during the same phase of the reproductive cycle. In this particular case, each month lasts 53 days and the raw NDVI values, originally spaced of 16 days, are spaced of 28 days.

Lastly, to investigate the influence of infant mortality (H6), we modified the proportion of infants dying from external causes by a coefficient *m* (see submodels “Death” in supplementary for details). The coefficient *m* took three different values, 0, 1 and 4, corresponding respectively to 0, 11.61 and 46.44% of infants dying from external deaths and representing the range observed in baboon species (Palombit, 2003).

## Results

We created a model simulating the life cycle of a female yellow baboon. Under realistic conditions, we found as output of our model comparable key life-history traits and reproductive seasonality (non-seasonal reproduction) to the ones observed in the wild, thus confirming its validity (Appendix C, Table S5 and Figure S5). We then modified environmental and life history parameters to test our hypotheses (H1-6).

## 1. Substantial support for the ecological hypotheses (H1-3)

An increasing gradient of birth seasonality was observed for increasing values of environmental seasonality and decreasing values of environmental productivity (Fig. 2a, top-left to bottom-right diagonal). High birth seasonality was therefore observed for environments characterised by both a high seasonality and a low productivity that still sustained a viable reproduction (non-grey cells in Fig. 2a). In very seasonal and very unproductive environments (bottom right corner of the heatmap), baboons’ fitness was indeed of zero, regardless of the phenology strategy followed (Fig. S6). Birth seasonality increased with environmental seasonality but this effect almost disappeared when environmental productivity was high (Fig. 2b). Similarly, birth seasonality decreased when environmental productivity increased, and this effect was modulated by environmental seasonality: the higher the environmental seasonality, the higher the effect of environmental productivity on birth seasonality (Fig. 2c). More precisely, high birth seasonality was observed only for the lowest values of environmental productivity.

**Figure 2:**
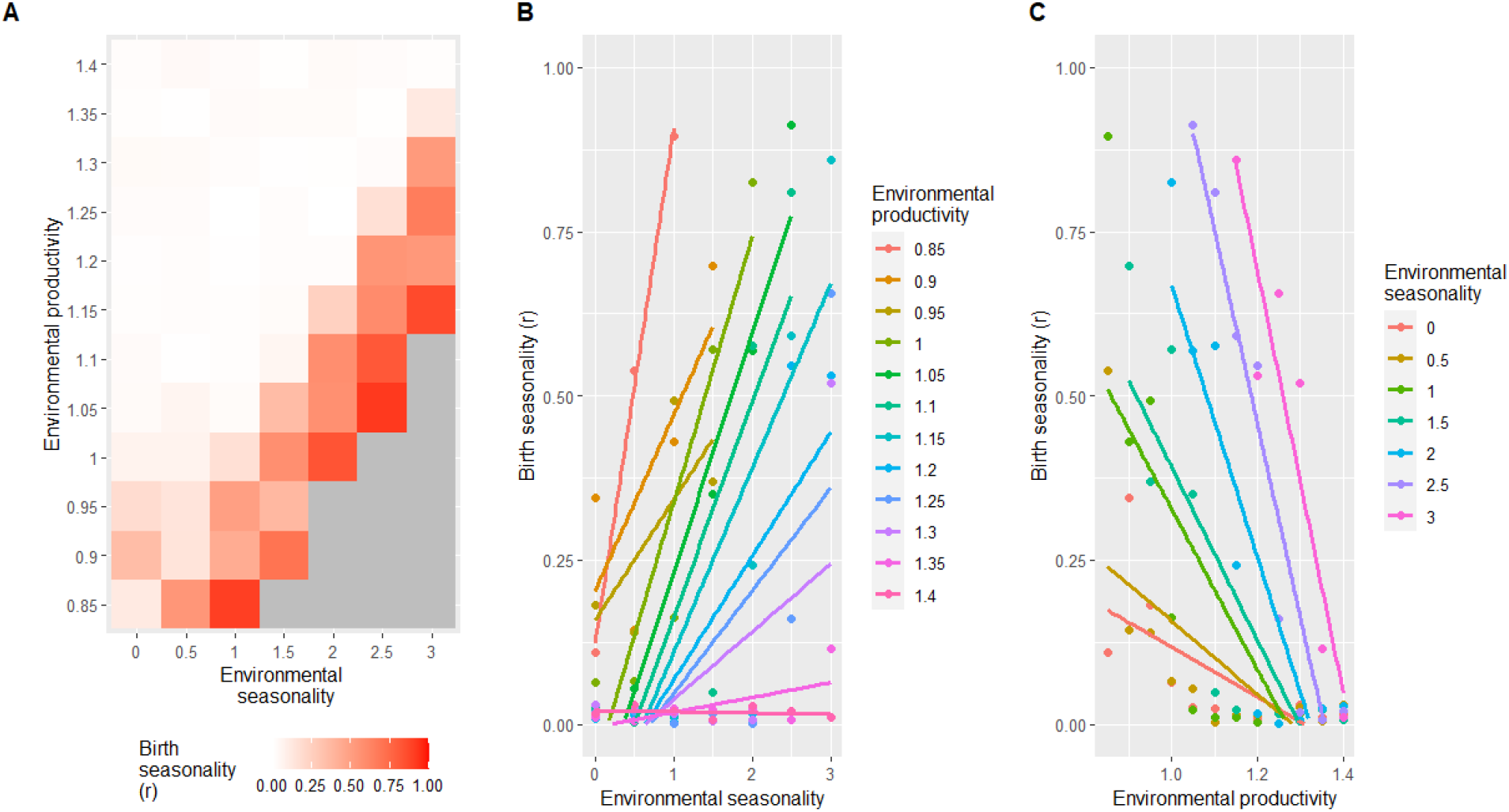
Effect of environmental seasonality and productivity on birth seasonality (H1-2) Panel A shows a heatmap of birth seasonality along gradients of environmental seasonality and productivity (normal conditions are represented by a seasonality and a productivity of 1). The other parameters remain unchanged (unpredictability of 1, growth rate of 5g/day, IBI of 1.7 years, infant mortality of 11.61%). Birth seasonality is represented by *r*, going from 0 (births are equally distributed) to 1 (all births occur on the same day). Grey cells represent non-viable environments characterised by a fitness of zero (regardless of the phenology strategy followed), where r cannot be computed and generates missing values. Panel B and C represent the same data, focusing on the effect of environmental seasonality for panel B and productivity for panel C, in interaction with the other.

The variation of birth seasonality observed throughout this environmental landscape results from various levels of selection pressure (Fig. 3a-c), with a steeper selection in seasonal environments leading to highest birth seasonality (Fig. 3c).

**Figure 3:**
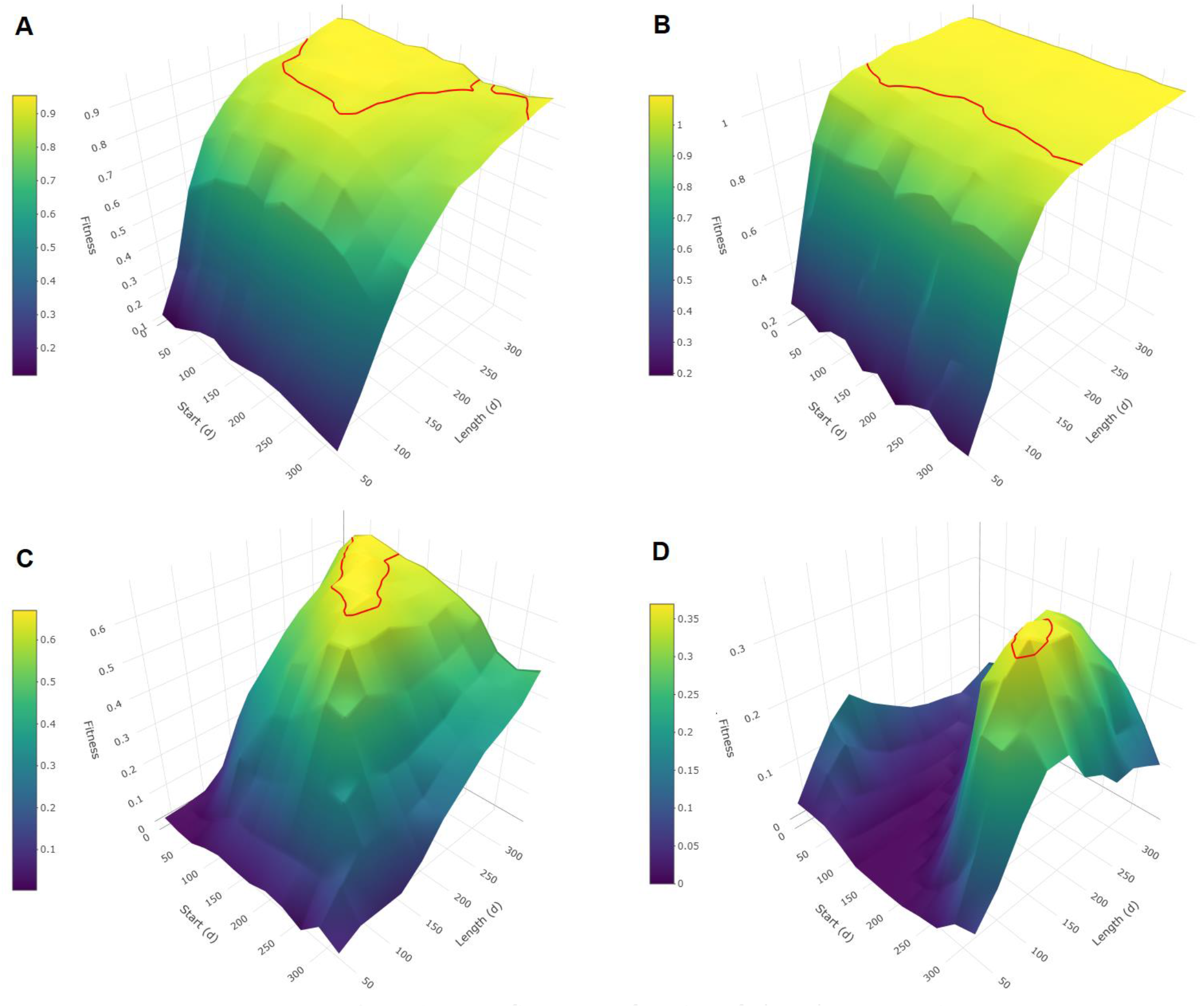
Mean fitness as a function of phenology strategies. Each 3D surface plot represents mean fitness (individual population growth rate, λind) as a function of the phenology strategy followed, for different sets of parameters. For each phenology strategy, mean fitness is obtained from 2000 simulations of the life cycle of a female following this strategy. Phenology strategies are described by a time window with a specific start (day of year) and a specific length (in days) when the female is allowed to conceive. The red line delineates the optimal phenology strategies, defined as strategies whose mean fitness value ranges above 95% of the highest mean fitness value. The steepness of the slope around these strategies reflects the intensity of the selection pressure towards them, with a flat curve corresponding to a weak selection pressure. In panel A, environment and life history correspond to the realistic conditions in Amboseli. In each of the other panels, only one simulation parameter is altered from the realistic conditions. Panel B shows a more productive environment (productivity = 1.1) and panel C shows a more seasonal environment (seasonality = 1.5), while environmental unpredictability and life history remain unchanged. Panel D shows an increased daily reproductive energy expenditure (growth rate = 7.5 g/day) while the environmental parameters and the other life-history parameters remain unchanged.

Modifying environmental unpredictability (Fig. 4) did not have an effect on mean birth seasonality (*r_mean_* = 0.17, 0.20, 0.18 and 0.19 for *u* = 0, 1, 2 and 3 respectively). However, it had an effect on the variance of birth seasonality values observed within each heatmap (*sd* = 0.28, 0.28, 0.19 and 0.11 for *u* = 0, 1, 2 and 3 respectively) with a strong decrease in maximal values (*r_max_* = 0.96, 0.91, 0.64 and 0.38 for *u* = 0, 1, 2 and 3 respectively). Yet, this effect was observed only for high values of unpredictability (*u* = 2 or 3), and we did not observe any effect on birth seasonality when completely removing the unpredictability of the environment (*u* = 0). Finally, here again, the effects of environmental traits on reproductive seasonality were modulated by each other: the higher the environmental unpredictability, the weaker the effects of environmental productivity and seasonality on birth seasonality.

**Figure 4:**
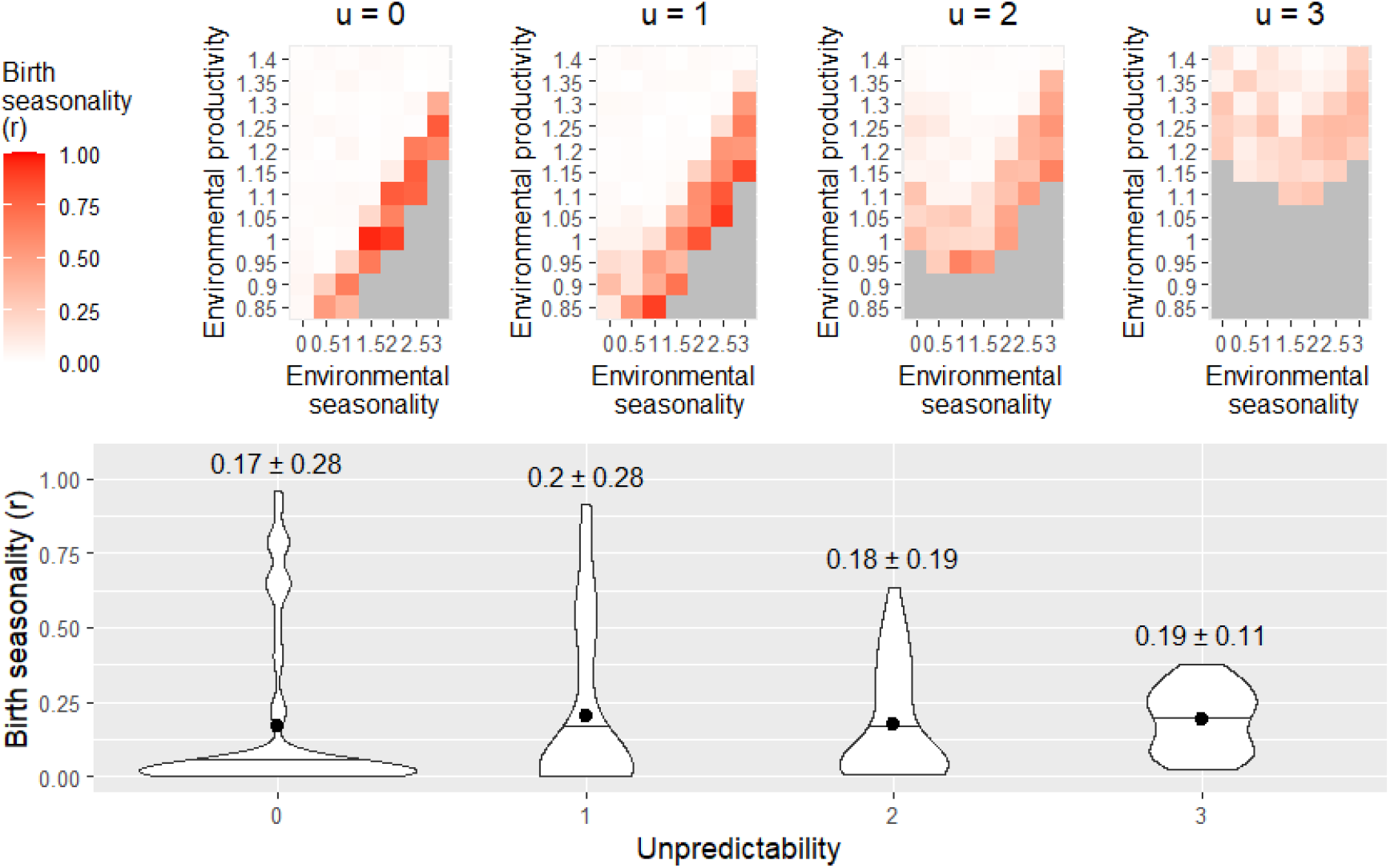
Effect of environmental unpredictability on birth seasonality (H3) The top panel shows heatmaps of birth seasonality along gradients of environmental seasonality and productivity in four different conditions of environmental unpredictability: no unpredictability (*u* = 0), normal unpredictability (*u* = 1) and increased unpredictability (*u* = 3 and 4, respectively). The other parameters remain unchanged (growth rate of 5g/day, IBI of 1.7 years, infant mortality of 11.61%). Birth seasonality is represented by *r*, going from 0 (births are equally distributed) to 1 (all births occur on the same day). Grey cells represent non-viable environments characterised by a fitness of zero (regardless of the phenology strategy followed), where r cannot be computed and generates missing values. The panel below shows the distribution of birth seasonality values observed in each heatmap. Black dots indicate mean values while black horizontal lines indicate median values. Means and standard deviations are given above each distribution (mean ± sd).’

## 2. Strong support for the daily reproductive energy expenditure hypothesis (H4)

Increasing the daily energy expenditure during reproduction by increasing the daily growth rate from 5 g/day to 7.5 g/day (+50%) dramatically increased the birth seasonality (+165%) even in environments characterised by high environmental productivity or low environmental seasonality (Fig. 5, *r_mean_* = 0.20 and 0.53 for growth rates of 5 g/day and 7.5 g/day, respectively). This increase in birth seasonality with daily energy expenditure was associated with a steeper selection pressure (Fig. 3d). Individual fitness simultaneously decreased for practically all environmental conditions (Fig. S6).

**Figure 5:**
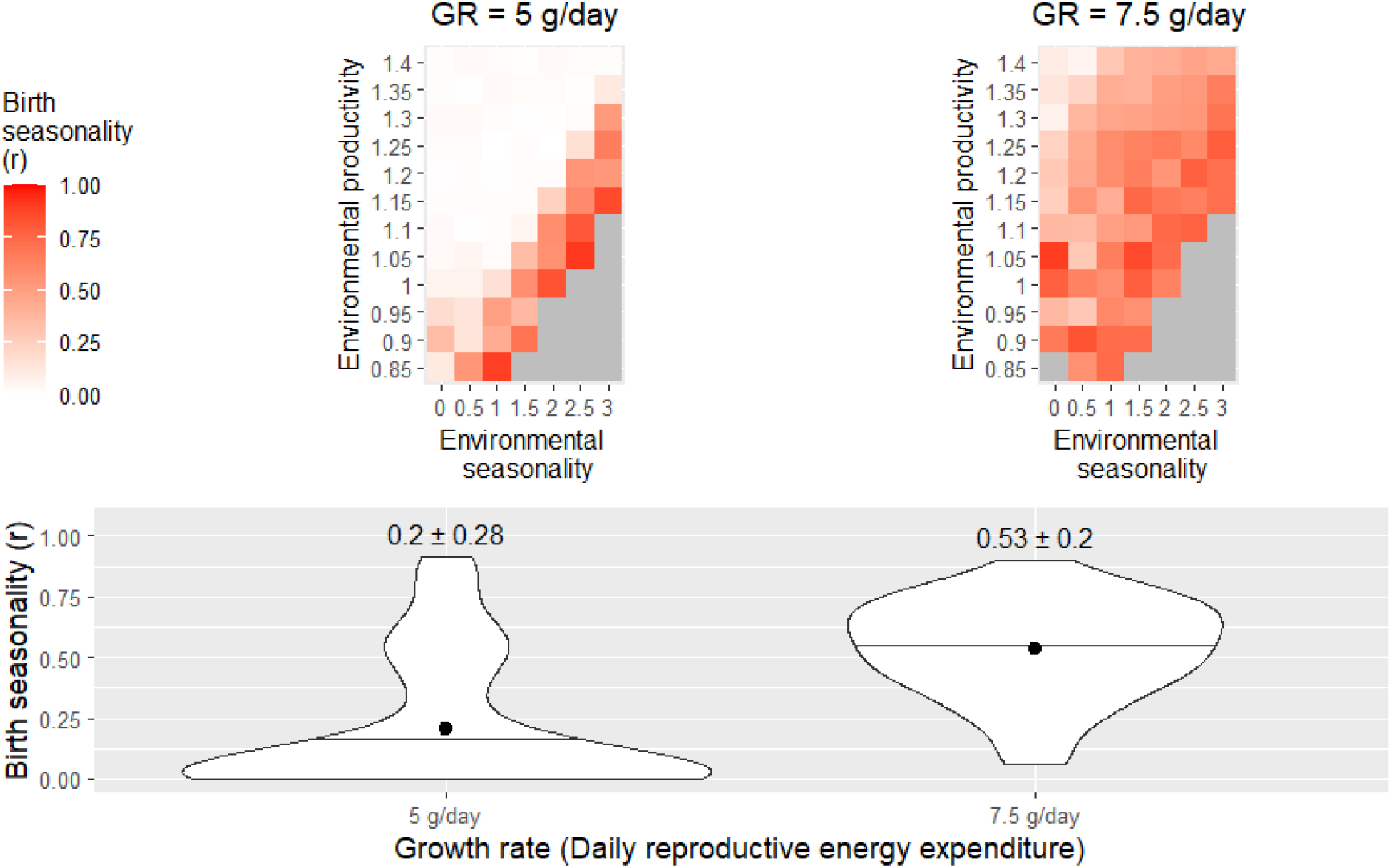
Effect of daily reproductive energy expenditure on birth seasonality (H4) The top panel shows heatmaps of birth seasonality along gradients of environmental seasonality and productivity in two different conditions of daily reproductive energy expenditure: regular (growth rate = 5 g/day) and increased (growth rate = 7.5 g/day). The other parameters remain unchanged (unpredictability of 1, IBI of 1.7 years, infant mortality of 11.61%). Birth seasonality is represented by *r*, going from 0 (births are equally distributed) to 1 (all births occur on the same day). Grey cells represent non-viable environments characterised by a fitness of zero (regardless of the phenology strategy followed), where r cannot be computed and generates missing values. The panel below shows the distributions of birth seasonality values observed in each heatmap. Black dots indicate mean values while black horizontal lines indicate median values. Means and standard deviations are given above each distribution (mean ± sd).

## 3. Minimal support for the length of reproductive cycle hypothesis and no support for the infant mortality hypothesis (H5-6)

No strong effect of reproductive cycle length on birth seasonality was observed (Fig. 6a and S7a), although we detected a tendency suggesting that birth seasonality was higher when interbirth intervals lasted exactly one year (*r_mean_* = 0.28, 0.23 and 0.20 when IBI lasted 1, 1.5 and 1.7 years, respectively).

**Figure 6:**
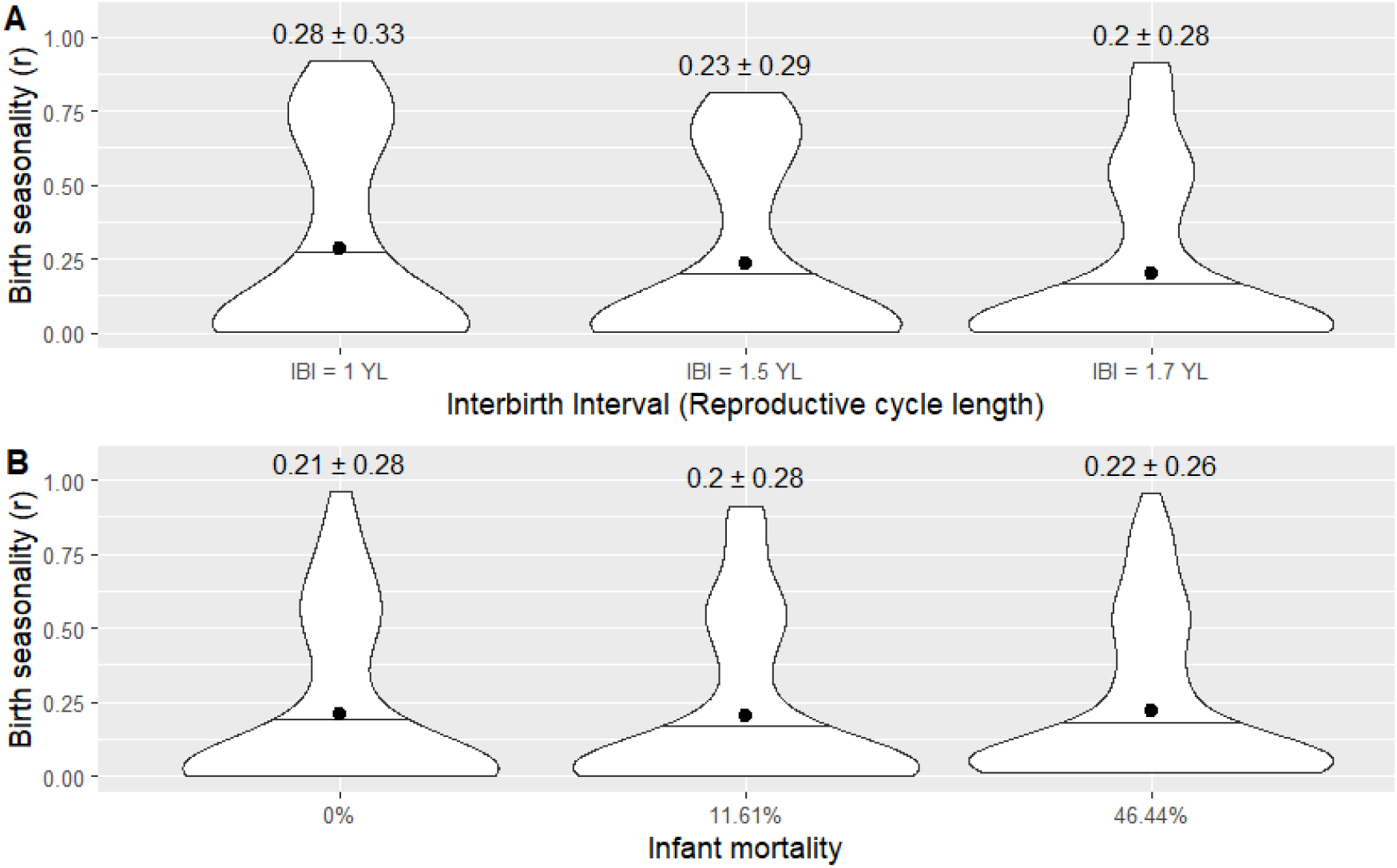
Effect of reproductive cycle length and infant extrinsic mortality on birth seasonality (H5-6) Panel A shows distributions of birth seasonality along gradients of environmental seasonality and productivity in three different conditions of reproductive cycle length: when the interbirth interval (IBI) is exactly one year long (YL), when its length is 1.5 years, and in normal conditions, when it is 1.7 years long. The other parameters remain unchanged (unpredictability of 1, growth rate of 5g/day, infant mortality of 11.61%). In panel B, we plotted distributions of birth seasonality along gradients of environmental seasonality and productivity in three different conditions of infant extrinsic mortality rate (M), respectively of 0%, 11.61% (observed rate) and 46.44%. The other parameters remain unchanged (unpredictability of 1, growth rate of 5g/day, IBI of 1.7 years). The associated heatmaps (where the plotted distributions come from) are presented in supplementary materials (Fig. S7). Birth seasonality is represented by *r*, the length of the mean vector, going from 0 (births are equally distributed) to 1 (all births occur on the same day). Black dots indicate mean values while black horizontal lines indicate median values. Means and standard deviations are given above each distribution (mean ± sd).

Birth seasonality did not increase after cancelling infant extrinsic mortality nor decreased when infant mortality increased, despite the substantial effect sizes that we explored regarding mortality rates (four-fold increase) (Fig. 6B and S7B, *r_mean_* = 0.21, 0.20 and 0.22 for m=0%, 11.61% and 46.44%, respectively).

## Discussion

### 1. Effect of ecology on reproductive seasonality

Unsurprisingly, ecology appears to play a key role in shaping reproductive seasonality. As expected, high levels of birth seasonality were associated with high environmental (H1). Yet, this effect gets weaker when the level of environmental productivity increases (H2). These two effects are in line with many studies showing a positive correlation between latitude and reproductive seasonality in most mammalian orders (Di Bitetti & Janson, 2000; Janson & Verdolin, 2005; English *et al*., 2012; Zerbe *et al*., 2012; Heldstab *et al*., 2018; Heldstab *et al*., 2021; Heldstab, 2021b, 2021a). While this association is often interpreted as the effect of environmental seasonality alone, increasing latitude reflects both increasing environmental seasonality and decreasing environmental productivity. Yet, the effect of environmental productivity has been overlooked in previous studies, with mean temperature being the closest proxy found to have a negative effect on reproductive seasonality, even when corrected for latitude (Heldstab, 2021b, 2021a; Heldstab *et al*., 2021).

In addition, our results support H3, by showing that high environmental unpredictability promotes non-seasonal breeding strategies while controlling for environmental seasonality and productivity; this hypothesis has often been proposed but rarely tested. These results are of major interest in the context of climate change, which increases the frequency and severity of extreme events (Easterling *et al*., 2000). Environmental unpredictability could even be a third overlooked factor encapsulated in the latitude effect, even if its relationship with latitude is not straightforward (English *et al*., 2012; Botero *et al*., 2014). In line with this, English *et al*. (2012) showed that birth seasonality in ungulates was better explained by the interaction between NDVI constancy and contingency – two components of predictability (Colwell, 1974) – than by latitude alone. However, another recent study investigating the hypothesis that non-seasonal breeding in chacma baboons was linked to environmental unpredictability found no support for it (Dezeure *et al*., 2023). These mixed results call for more empirical studies, which should disentangle environmental productivity, seasonality and unpredictability in order to reach firm conclusions.

The choice of proxy used to describe environmental unpredictability is most likely critical to explain the mixed results observed for the effect of unpredictability. Here, we use between-year variation as a proxy for environmental unpredictability, while this component of environmental variation could be at least in part predictable (e.g. El Niño climate events, Chen *et al*., 2004). Additionally, in this study we focused on NDVI variation in *magnitude*, while variation in the *timing* of the annual peak of NDVI could also be critical. For example, environmental conditions characterised by a food peak whose timing differs between years – while still occurring within the same annual period of 3-4 months - would be captured by the non-seasonal component of NDVI, even though the majority of higher-than-average values occur around the same time of the year, during the (variable) seasonal food peak. This implies that this non-seasonal component of NDVI encapsulates, in fact, some part of seasonality in its timing and may explain why, in our model, removing environmental seasonality (*s* = 0) is not enough to observe complete non-seasonal reproduction. To do so, we further need to remove environmental unpredictability (*u* = 0). In this context, our methodological approach relying on the NDVI decomposition to disentangle the effect of environmental seasonality and unpredictability may therefore not be sufficient, and future studies would usefully find a way to distinguish unpredictability in the magnitude and in the timing of the annual food peak, which may influence the ecology and evolution of seasonal breeding in different ways.

## 2. Effect of life-history on reproductive seasonality

Low day-to-day reproductive energy requirements proved critical in enabling non-seasonal reproduction (H4). Even a rather moderate increase in growth rate (×1.5), for a constant body mass, was sufficient to increase significantly birth seasonality regardless of environmental seasonality and productivity. This acceleration of reproductive pace has two distinct, and important consequences: (1) maximal daily reproductive energy needs increase for the mother, and (2) the most demanding stages of the reproductive cycle are circumscribed in a shorter period of time - which may only represent one fragment of the year cycle. These two changes, in combination, can lead to a situation where females can only sustain reproduction during the good season. Such an increase in reproductive pace – and daily reproductive costs – corresponds to a slight move on the slow-fast continuum (Bielby *et al*., 2007) that could be observed within phylogenetically close (and potentially sympatric) species and partly explain the diversity of reproductive seasonality observed among them. This could for example explain why mandrills (*Mandrillus sphinx*), despite being closely related to yellow baboons and living in the equatorial forests of the Congo basin, reproduce seasonally (Hongo *et al*., 2016; Dezeure *et al*., 2022). Growth rate in young mandrills is indeed higher (×1.34), and lactation periods are accordingly shorter than in yellow baboons (about half as long), even though gestation length, birth mass and weaning mass are comparable and adult female mandrills body mass is much lower than for female baboons (Setchell *et al*., 2001; Dezeure *et al*., 2022), meaning an even larger difference in daily reproductive energetic expenditure when controlled by maternal size for these two species. At the extreme end of life-history paces, are Bornean orangutans (*Pongo pygmaeus wurmbii*) which flattened their reproductive energy requirements so much that they reached a low and constant level of maternal effort (van Noordwijk *et al*., 2013), with nearly six years of lactation (compared to about one year in yellow baboons), which makes it easy to understand why birth timing does not matter much.

Notably, in our study model, an increase in growth rate, with a constant body mass and especially with a constant weaning body mass threshold, translates into an acceleration of the reproductive cycle, which could favour the evolution of seasonal breeding through another mechanism tested by hypothesis H5, namely that the reproductive cycle is closer to exactly one year, leading to the reduction of gaps between the successive reproductive cycles of a female, who must wait for the next reproductive season to start a new cycle after weaning an infant. However, when manipulating the duration of the reproductive cycle relatively to the annual seasonal cycle, we did not find any significant effect, confirming that the observed effect of growth rate is not caused by a change in the length of the mother’s reproductive cycle as expected under H5, but indeed by the increase of her daily reproductive energy expenditure itself as expected under H4.

Even though this effect of daily reproductive energy expenditure on the evolution of reproductive seasonality has, to our knowledge, never been formally tested, our results are corroborated by some empirical studies. For example, a shorter gestation was found to be associated (when controlling for body mass) with a higher reproductive seasonality in several mammalian orders (Zerbe *et al*., 2012; Heldstab *et al*., 2018; Heldstab *et al*., 2021) ; such a result could, in fact, reflect the increase in daily reproductive energy requirements due to a shortened gestation. This interpretation is strengthened by the case of lagomorphs where the intensity of reproductive seasonality was not associated with gestation length but with litter size (Heldstab, 2021a), with an increased breeding seasonality for species with larger litters, facing increased daily reproductive energy requirements. Overall, these comparative studies that reported associations between reproductive seasonality and various aspects of reproductive pace are consistent with our results supporting a single life-history hypothesis, namely that daily reproductive energy intake is a major determinant of the evolution of (non)seasonal breeding.

Our last hypothesis stated that non-seasonal reproduction would emerge from a high mortality rate of infants compared to adults (H6), but we could only detect a weak, tendency in support of H6. It was already known that high rates of infanticide (leading to a high mortality rate of infants relatively to adults) are associated with non-seasonal breeding (Lukas & Huchard, 2014). It is commonly admitted that this is because infanticide is only beneficial for male killers when it accelerates female’s return to cycle substantially, which may rarely occur in seasonal breeders where females only re-cycle in the next mating season. Even if our results do not support the alternative interpretation where infanticide could instead select for non-seasonal breeding, the tendency we detected could suggest a co-evolutionary pattern where both traits reinforce each other.

## 3. Insights on the emergence of reproductive seasonality

High reproductive seasonality (over 0.75) was almost systematically associated with low individual fitness values (under 0.25) in our simulations. Even though such simulated measures of individual fitness cannot accurately inform about demographic dynamics, very low mean values of individual fitness may represent ecological situations with declining populations, where reproductive seasonality is unlikely to evolve successfully, at least in theory. Yet, such theoretical considerations are mitigated when confronted to empirical figures: in our simulations, moderate reproductive seasonality (around 0.5, which corresponds to the highest value observed within the genus *Papio*, Petersdorf *et al*., 2019) is often associated with mean values of individual fitness that are comparable to those calculated from the Amboseli baboons (λ_ind_ = 0.9, Fig. S6, tile *s*=1 and *p*=1 in the top left heatmap), where the long-term demographic trend seems stable (Samuels & Altmann, 1991), so that we cannot rule out the possibility for seasonal reproduction to emerge on the basis of low values of individual fitness.

In addition, it’s important to underline some limitations of our modelling strategy. In particular, sociality could play an additional role and modulate the effects of environment and life history examined here. For example, in some prey species, giving birth synchronously may decrease predation risk via predator satiation (Ims, 1990). In other cases, synchronous births may instead enhance reproductive competition (over food, access to mates or paternal care), and increase the costs of seasonal reproduction. In line with this, it has recently been showed, in a wild chacma baboon population (*Papio ursinus*), that the reproductive timings of subordinate females were influenced by the reproductive state of other females of the group, leading to a lower reproductive synchrony in the group, thus decreasing breeding seasonality at the population level (Dezeure *et al*., 2023). Such a polymorphism of strategies, where some females may choose to delay their reproduction or move it forward to minimize competition over limited resources, could not emerge from our model that only optimises the strategy of an isolated female in the absence of competitors. In order to integrate such frequency-dependent effects, other models, such as game theoretic models, should be developed. Additionally, and on top of sociality, dynamic individual strategies are not considered in this study: a female’s investment does not vary dynamically as a function of her age and the model does not allow for terminal investment or increase in reproductive investment toward the end of life. Such variation could be best modelled with dynamic state modelling (dynamic programming).

## 4. Generalisation potential

The general theoretical framework developed in this study and in particular most of our working hypotheses could apply to most species experiencing substantial reproductive costs, except perhaps for H6, the hypothesis on infant mortality rates, which is restricted to species with small litters or singletons that can be lost all at once. As such, the ecological and life-history factors that were identified as influential in our model could, potentially, have a wide predictive power. Nevertheless, because our model was deemed realistic, we only tested our hypotheses in local conditions, meaning that our results may not be valid across all environmental conditions, or across the entire life-history space. In particular, adapting our model to a faster-living species, or running a comparative analysis testing these hypotheses across a large taxonomic group encompassing variable environments and life-histories, could bring valuable insights on the generalisability of our results.

Meanwhile, these results can reasonably be extended to other slow-living species, especially slow-living primates. As such, they shed new light on the emergence and maintenance of non-seasonal breeding in apes and humans (Bronson, 1995). As other slow-living primates, humans experience low daily maternal energy costs spread over multiple years (Dufour & Sauther, 2002), and this trait may itself be sufficient to explain why they breed non-seasonally, as they otherwise live in an exceptionally wide range of environmental conditions. Other characteristics that have often been evoked in this context, and are sometimes seen as human-specifics, such as cumulative culture and associated niche construction, agriculture and technology, communal breeding or extreme behavioural flexibility (Bronson, 1995) may, in fact, not be required to explain why humans breed year-round. However, these traits likely provide humans with additional capacity to extract energy from harsh and fluctuating ecological conditions, possibly allowing them to sustain faster reproductive pace, and strive in a wider variety of environments, than other large primates (Wells & Stock, 2007).

## Conclusion

This study contributes to the development of a broad theoretical framework on the evolution of reproductive seasonality, which, despite being a classical topic in evolutionary sciences, remains lacunary. We tested this framework through a set of hypotheses, by constructing an original and realistic agent-based model parameterized on a slow-living primate. We could show that ecology and life history interact to shape the emergence of reproductive seasonality. Specifically, this study disentangles the effect of three main components of environmental variation, namely seasonality, productivity and unpredictability, which probably all contribute to generate the well-known association between latitude and reproductive seasonality. We further highlighted how high daily reproductive energy requirements - which typically translate into a fast life history pace – strengthen the intensity of reproductive seasonality, independently of environmental variations. Overall, this study contributes a powerful theoretical framework and modelling tool that may apply across the life-history space, as well as sheds new light on the unexplained variance in the reproductive phenology of long-lived species including humans.

## Acknowledgements

The results of this study were partly produced with the support of the Montpellier Bioinformatics Biodiversity MBB platform. We would like to thank Jimmy Lopez and Khalid Belkhir for their assistance in structuring the model and using the MBB platform.

## Data, scripts and codes availability

Program code used in this study are freely available in the GitHub repository https://github.com/lburtschell/RepSeason

## Supplementary material

Supplementary material of this study is available in the GitHub repository https://github.com/lburtschell/RepSeason

## Funding

This work was supported by the ANR grant ANR ERS-17-CE02-0008, 2018-2021 awarded to Elise Huchard. Lugdiwine Burtschell thanks AgroParisTech for the financial support.

## Notes

https://github.com/lburtschell/RepSeason

### Competing Interest Statement

The authors have declared no competing interest.

### Summary of Updates

Recommendation of this article has now been published by PCIEcology with the editorial correspondence. We added the PCIEcology badge on the 1st page of the article with a link to the recommendation.

