## Supplementary material for "Evolutionary determinants of reproductive seasonality: a theoretical approach"

#### APPENDICES

##### *Appendix A. Description of agents' state variables*

Individuals are described by state variables, presented in Table S2. The first state variable is the total mass (in kilograms) of the individual, divided in two compartments, lean mass and fat mass. Lean mass increases with the growth of the individual while fat mass describes its energy reserves. Two other attributes (namely "Alive" and "Growth Allocation" in Table S2) describe if individuals are alive and how they allocate their energy to growth (depending on the resources they get). Each individual is also characterised by the timing of key life events: cycling, conception, birth, weaning, sexual maturity, death (Fig. 1). These timings are expressed in days, and 0 corresponds to the first day of the simulation. A lifespan is also assigned to each individual and represents their maximal life duration. It can equate the longevity of the species or be lower to account for extrinsic mortality (e.g. predation, disease, infanticide). For foetuses, this attribute represents the gestation duration, either equal to the usual gestation length or lower in case of miscarriages. Individuals can also die before this lifespan is reached if they fail to find sufficient energy supplies at any point of the simulation. Energetically speaking, we compute the energy balance of each individual who has reached nutritional independence (juvenile or female) by comparing its needs to the available energy. Energy needs represent the amount of energy needed for maintenance, growth and (in adults only) reproduction while available energy is the sum of the energy intake (extracted from the environment) and the energy that can be released from storage. In addition, each female follows a given phenology strategy that remains constant throughout the simulation and which sets the starting point and length of a time window during which she can cycle and conceive. For each reproductive cycle, females are attributed a cycling duration (which can vary) before they conceive. Females are also characterised by their current reproductive status: anoestrus, oestrus, gestation or lactation (Fig. 1) which is associated with the list of their offspring at each life stage (foetus, infant and juvenile). The values of the initial attributes for each life stages are presented in table S3 and S4.

***Appendix B. Submodels***

#### 1. Death

An individual dies if it reaches its assigned lifespan (external mortality causes or natural death) or if his lean mass falls beyond a critical threshold (starvation). If a female dies, her current foetus or infant dies as well, but juveniles survive because they are nutritionally independent from their mother.

*1.1 External mortality:*

To account for external mortality causes or natural deaths in females, we used data of observed breeding female lifespans in the Amboseli population of yellow baboons (McLean *et al.*, 2019) to assign randomly a realistic lifespan value to each adult female at the beginning of each simulation (96 values ranging from 2234 days or 6.1 years to 9457 days or 25.9 years). We used the same process to account for external mortality (or miscarriage) in the other life stages: foetuses, infants and juveniles.

In the wild, 13.9% of pregnancies result in foetal losses (Beehner *et al.*, 2006). One proportion of those miscarriages, due to insufficient maternal energetic intake, was already taken into account by the model (see submodel "Energy allocation"), but we needed to add an external probability of miscarriage to account for other causes of miscarriages. Because the actual value is unknown, we chose a probability so that the overall miscarriage rate in the model fits the 13.9% observed in the wild as closely as possible. This external proportion of miscarriages was 7%. As a result, in the model, 7% of foetuses were assigned a lifespan between 1 and 178 days (the average gestation length), meaning that they died before birth.

The proportion of infants dying from external causes before weaning was fixed at 54% of the overall mortality, corresponding to the proportion of infant deaths suspected or confirmed to be violent deaths (Alberts, 2019). The overall mortality between birth and weaning in a natural population being of 21.5% (McLean *et al.*, 2019), we estimated the proportion of infants dead from external causes at 21.5% * 54% = 11.6%. Consequently, 11.6% of infants were assigned, at birth, a lifespan that was randomly picked from those infant lifespans observed in the Amboseli population which were inferior to the average weaning age (116 values ranging from 1 to 319 days). Other, surviving, infants were assigned a lifespan fixed at yellow baboon longevity (27 years or 9862 days, (Bronikowski *et al.*, 2002)).

Because there were no data on the causes of death for juveniles, we used the same external mortality rate as for infants, assuming that the low infanticide rate in Amboseli, responsible for external deaths of infants but not juveniles, was insufficient to justify a diminishing of the external mortality for juveniles, especially considering that the overall mortality for juveniles (from weaning to sexual maturity) is slightly higher than for infants (McLean *et al.*, 2019). 11.6% of juveniles were therefore also assigned, at weaning, a lifespan that was randomly picked from juvenile lifespans observed in the Amboseli population which corresponded to death after weaning and before sexual maturity (128 values ranging from 329 to 1731 days).

*1.2 Starvation:*

Individuals die from starvation when they take too much energy from their lean mass, after having used up all the energy available in the environment and in their fat reserves. We estimated the critical threshold for lean mass from two different studies. The first one calculates the energy requirement for growth and estimates that healthy tissue is composed at 84% of lean mass (Payne & Waterlow, 1971). The second one is a study on women with anorexia (Scalfi *et al.*, 2002) whose lean mass when malnourished represented 86% of their lean mass after complete recovery. Thus, we estimated in the model that when lean mass falls below 85% of its regular value, the individual would die.

#### 2. Change of reproductive status

The female reproductive cycle consists in four successive phases: Oestrus, Gestation, Lactation and Anoestrus, each occurring after defined events: cycling, conception, birth and weaning (Fig.1).

*2.1 Cycling:*

There are two conditions for a female to switch from the "anoestrus" to the "oestrus" status. First, the current time t must be within the time window defined by the female phenology strategy. Second, as showed in Gesquiere *et al.*, 2018, the female needs to have a positive energy balance, with an increase in body fat for at least one month. If the female is still cycling (i.e. has failed to conceive) when the end of the reproductive window is reached, she goes back to "anoestrus".

*2.2 Conception:*

To enter the "gestation" phase, a female has to conceive. Gesquiere *et al.*, 2018 showed that duration of cycling until conception is highly variable (138 ± 82 days, cv = 60%) but insensitive to environmental variation. We thus chose in the model to randomly pick this duration from observed data in Amboseli (Gesquiere *et al.*, 2018) at the beginning of each oestrus period.

Once gestation starts, a foetus is created (Table S4) and added to the female "list of foetuses" attribute. There is always a single foetus produced (Altmann, 1983).

*2.3 Birth:*

Birth occurs after a constant gestation length of 178 days (Gesquiere *et al.*, 2018). The female's reproductive status changes then to "lactation", an infant is created using the foetus attributes (Table S4) and added to the female "list of infants" attribute.

*2.4 Weaning:*

Weaning is a continuous process whose beginning and end are difficult to quantify. Cessation of suckling and complete feeding independence is reached at approximately 595 days (85 weeks) of age in yellow baboons (Altmann, 1998). Nevertheless, the observed lactational anoestrus is much shorter, around 322 days (Gesquiere *et al.*, 2018) with resumption of cycling that corresponds to a drop in suckling time (Lee *et al.*, 1991). Moreover, it has been observed in yellow baboons that independence from the mother for nourishment and transportation was almost reached by 365 days of age (Rhine *et al.*, 1985), even if occasional suckling often continues throughout gestation (Lee *et al.*, 1991). To simplify the model, we chose to assume a complete weaning around the age of 46 weeks (322 days), corresponding to the mean post-partum amenorrhea duration in Amboseli (Gesquiere *et al.*, 2018). Nevertheless, it has been shown that weaning is better explained by a threshold mass than a threshold age (Lee, 1996). Because total mass can vary between same-age individuals, this condition allows us to introduce variability in the postpartum amenorrhea duration, as observed in wild populations (Gesquiere *et al.*, 2018). We thus calculated the weaning threshold total mass at 2.32 kg (birthMass + ppaDuration x growthRate). After weaning, the infant becomes a juvenile (Table S4) and the mother goes back to the anoestrus status. During gestation or lactation, if the foetus or the infant dies, the female goes back to anoestrus.

#### 3. Energy intake

Females and juveniles feed every day. Food availability is approximated by NDVI, and we assume that saturation occurs for high NDVI values, while for low NDVI values, alternative sources of food, called “fallback foods”, (Altmann, 1998; Brockman & van Schaik, 2005) are used. Therefore, we chose to describe energy intake as a saturating function of NDVI (Fig. S1) as follows:

$$E(m,t) = E_{max}(m) \times \left(1 - exp^{-\frac{NDVI(t)}{\tau}}\right) \text{ (Equation 1)}$$

Where $E_{max}$ is the maximum value of energy intake for a baboon of mass *m*, when food is non-limiting and $\tau$ is the characteristic value of the equation, representing the speed at which $E_{max}$ is reached, and therefore the efficiency of fallback foods. The parameter $\tau$ being unknown, we used it to calibrate our model. Technically, we tried various values until obtaining realistic life history traits in the model outputs (Table 1, main text) and found the value of $\tau = 0.117$.

We used a study calculating energy intake for non-reproductive adult female baboons (*Papio cynocephalus* and *Papio anubis*) fed *at libitum* to estimate $E_{max}$ at 4891.1 kJ/day (Roberts *et al.*, 1985) for adult females of $m_{adult}$ =11.9 kg (Altmann *et al.*, 1993). Because energy needs increase allometrically (Kleiber, 1932), we chose to reproduce this variation for energy needs, with $E_{max}$ being a function of mass *m* to the power 0.75 (see submodel "energy needs" for details):

$$E_{max}(m) = 4891.1 \times \frac{m}{m_{adult}}^{0.75} \quad \text{(Equation 2)}$$

Similarly, a female's energy needs increase during reproduction, and so does her energy intake. For gestating females, this increase in energy intake is considered through an increase in total mass (Equation 2). For lactating females, Roberts *et al.*, 1985 reported that energy intake increased by 11% at the beginning of lactation with a maximum of 27% at the beginning of weaning. We translated this increase in energy intake through a multiplying coefficient *c*, varying linearly with infant mass $m_i$ and therefore with lactation progress (Fig S2). In a first phase, when the infant is completely dependent, *c* increases from 1.11 at birth to a maximum of 1.27 at beginning of weaning:

$$c_1\ (m_i) = 1.11 + (1.27 - 1.11) \times \frac{m_i - birthMass}{massAtBeginningOfWeaning - birthMass} \quad \text{(Equation 3)}$$

In a second phase, when the infant starts to become independent, *c* decreases back from 1.27 to a final value of 1 when the infant is completely weaned and lactation stops:

$$c_2\ (m_i) = 1.27 + (1 - 1.27) \times \frac{infantMass - massAtBeginningOfWeaning}{weaningMass - \ massAtBeginningOfWeaning} \quad \text{(Equation 4)}$$

Moreover, it is reasonable to think that energy extraction efficiency changes with age, increasing rapidly at first and reaching an asymptote at last. We used again the same saturating function type and obtained a final equation for energy intake, depending on the time *t*, the mass of the female *m*, her age *a*, and the mass of her dependant infant $m_i$:

$$E_{intake}(t,m,a,m_i) = c(m_i) * 4891 \times \left(\frac{m}{m_{adult}}\right)^{0.75} \times \left(1 - exp^{-\frac{NDVI(t)}{\tau}}\right) \times \left(1 - exp^{-\frac{a}{\tau_{age}}}\right)$$

(Equation 5)

We chose $\tau_{age}$ in order to model a realistic offspring survival (Table 1, main text) and fixed it to $\tau_{age}$ = 193 days, giving a foraging efficiency of 81% for a juvenile just weaned (322 days).

With this final equation, we obtain, for the average NDVI in Amboseli (0.228), a daily energy intake for an adult female at the beginning of lactation of about 4670 kJ. This value is consistent with the average observed value in Amboseli for mixed groups of pregnant females, lactating females and males (4620 kJ/day for a 12kg baboon (Stacey, 1986)).

#### 4. Energy needs

Energy needs in the model consist in two main components: (i) energy for maintenance and travel ($E_m$), and (ii) energy for growth maintenance (*Eg*). The energy for reproduction can be seen as extra energy for maintenance, travel and growth of the offspring.

##### *4.1 Energy for maintenance and travel:*

Altmann & Samuels, 1992 estimated that a 11kg-female baboon alone, in Amboseli, would expend 3493.7 kJ/day for her own maintenance and travel. This value is the result of the sum of the cost of basal metabolism and energy spent for locomotion. The first one follows an allometric relationship to the power 0.75 of body mass (Kleiber, 1932) and the second one to the power 0.684 (Altmann, 1998). To simplify and because these two exponents are close, we chose to use 0.75 as the allometric exponent, which gives the following allometric relationship:

$$E_m(m) = A \times m^{0.75} \text{ (Equation 6)}$$

Based on the 3493.7 kJ/day spent by a 11kg female, we find that $A = \frac{3493.7}{11^{0.75}} = 578.41$, and we can re-write equation 6 as follow:

$$E_m(m) = 578.41 \times m^{0.75} \text{ (Equation 7)}$$

Where *m* is the body mass of a female, juvenile or infant, in kilograms. If the female is pregnant, *m* represents her total body mass (including foetus and placenta).

*4.2 Energy for growth:*

Growth in young baboons can be considered as linear with an average growth rate of 5g/day, indistinctly for males and females (Altmann & Alberts, 1987). We also assumed a linear growth during gestation, with a growth rate taking into account both foetus and placenta. We estimated the weight of the placenta to be 0.25 times the weight of the foetus (Farley *et al.*, 2009), which gives a growth rate during gestation of $\frac{1.25 \times birthMass}{gestationLength}$ = 0.005 kg/day. This simple calculation shows that growth rate is maintained between gestation and infancy. By simplicity we also kept this growth rate for juveniles (males or females) and adult females even though growth rate starts to increase and differ by sex after 4 years of age (Altmann & Alberts, 1987).

With a growth cost of 20.92 kJ/g (Payne & Waterlow, 1971), we can estimate energy needs for growth at all stages of life as:

$$E_g = 20.92 \times growthRate \text{ (Equation 8)}$$

We give to the individuals the possibility to slow or even pause their growth according to the resources they have, using their "growthAllocation" (*gA*) attribute, which equals 1 if growth is normal, 0 if growth is stopped and is between 0 and 1 if growth is slowed (see section Energy balance for detail). However, foetuses growth cannot be slowed, because pregnancy length varies very little (Gesquiere *et al.*, 2018). Except for gestation growth, equation 8 becomes:

$$E_g(gA) = gA \times 20.92 \times growthRate \text{ (Equation 9)}$$

*4.3 Total energy needs:*

For juveniles or non-reproductive adult females, total energy needs are the sum of energy needed for maintenance and travel and energy needed for growth:

$$E_N(m, gA) = E_m(m) + E_g(gA) \text{ (Equation 10)}$$

But during gestation and lactation, female energy needs are increased with additional needs for her foetus or infant. During gestation, the female covers all the energy needs of her foetus, and we neglect the costs associated to the energy transfer. Therefore, the total energy needs for a pregnant female, depending on her total mass (including foetus mass) and her growth allocation, are:

$$E_N^{pregnant}(m, gA) = E_m^{pregnant}(m) + E_g^{fem}(gA) + E_g^{foetus} \text{ (Equation 11)}$$

During lactation however, the energy transfer has a cost, due to the production of milk by the mother, which translates into a lactation efficiency of 0.8 (Dewey, 1997). We assume that there is no additional cost for the assimilation of milk by the offspring. In other words, if the infant needs 1 kJ of energy, the mother would have to give $\frac{1}{0.8} = 1.25$ kJ to cover such needs. Moreover, weaning being a continuous process, maternal lactation does not cover the full energy needs of an infant during lactation, but a decreasing fraction of it. We consider such decrease to be linear from when the infant reaches 0.99 kg (eighth week of life), that is, when he starts to significantly feed on plant food (Rhine *et al.*, 1985), and until complete weaning at 2.32 kg. Therefore, the total energy needs for a female during lactation are:

$$E_N^{lactating}\left(m, m_i, gA^f, gA^i\right) = E_m(m) + E_g^{fem}(gA^f) + \frac{1}{0.8} \times \alpha \times \left(E_m^{inf}(m_i) + E_g^{inf}\left(gA^i\right)\right)$$

(Equation 12)

$$where\ \alpha = 1\ if\ m_i \leq 0.99\ and\ \alpha = 1 - \frac{m_i - 0.99}{2.32 - 0.99}\ if\ m_i > 0.99$$

It should be noted that an infant with a mass over 0.99 kg has to cover a part of his own energy needs. Nevertheless, to simplify the model, we assume, without calculating it, that the infant succeeds at covering his remaining energetic needs, proportionally to the amount of energy its mother is able to give him (see submodel “Energy allocation” for detail).

#### 5. Energy allocation

A flowchart of energy allocation is presented in Figure S3. The first step is to compute the energy balance for the female. Energy balance is the difference between available energy and total energy needs. Energy needs represent the amount of energy needed for maintenance, growth and reproduction (see submodel “Energy needs”) while available energy is the sum of the energy intake (extracted from the environment, see submodel “Energy intake”) and the energy released from storage. On the one hand, if the energy balance is positive, the female can store the available energy in fat mass. One gram of fat mass contains 39.5 kJ (Livesey & Elia, 1988), but energy storage in the form of fat mass has an efficiency of 0.9 (Rothwell & Stock, 1982). If the female is lactating, she gives half of surplus energy to her infant. Because all ingested energy is not necessarily metabolised, we fixed a maximal daily amount of fat storage equal to the approximate amount of daily fat gain during growth. Baboons gain on average 5g per day of new tissue and have about 2% of fat in their body (Altmann & Alberts, 1987), which gives a maximal daily fat storage of 100 mg. Each day of the simulation, the extra energy that is not stored due to the storage limit is lost.

On the other hand, if the energy balance is negative, the female can reduce or even pause her growth for the day, depending on the amount of energy missing. If still necessary, she can reduce or pause her infant’s growth. This decision will reduce the amount of energy needed and could reverse the energy balance. If her energy balance is still negative after these savings steps, the female can try to use the available energy in her fat mass to cover her energetic costs. If after having burnt all her fat mass, her energetic needs are still uncovered, the female has no further choice than to abort if gestating, or abandon the infant if lactating. In both cases, the offspring dies and the female goes back to anoestrus. Finally, the last possibility for the female if her energy balance is still negative is to burn her lean mass, with 1g of lean mass releasing 20.92 kJ (Payne & Waterlow, 1971). If lean mass falls below a threshold of 85% of its initial value (before the female starts to burn lean mass), the female dies (see "death" submodel for details).

**6. Growth**

Once energy is allocated, the female and her potential infant or foetus can grow according to their "growth allocation" attribute. The "growth allocation" attribute can fall below 1 if growth is slowed, or even be null if there is not enough energy for growth, or if the female has reached her final adult size. The total daily body mass increase is therefore:

$$m_{increase} = growthAllocation \times growthRate \ (kg) \quad \text{(Equation 13)}$$

This daily increase is distributed between lean and fat mass according to the proportions observed in wild baboons (Altmann & Alberts, 1987) and an additional increase in fat mass can exist if the daily energy balance is positive (see submodel "Energy allocation").

We consider a fully grown female to be 11.9 kg (Altmann *et al.*, 1993), corresponding to a threshold lean mass of 11.6739 kg (adultMass x leanProportion). Yet, if an adult female has lost some of her lean mass because of a dearth of energy, she can grow again to rebuild these tissues. Because juveniles are independent from their mother, we estimate their growth by calculating their own energy balance (energy intake minus energy needs), with the same procedure as for a non-reproductive female.

#### *Appendix C. Model validation*

**1. Simulated values for key life history traits**

We conducted an internal validation by tracking every state variable throughout a simulation in search for any unrealistic value. We then checked the value of some key life history traits calculated from 2000 simulations and compared them to empirical values from the Amboseli wild population. In order to obtain relevant comparisons between simulated and observed values for each trait, we computed the simulated traits with the calculation methods described in each associated empirical study. Notably, and only in this part of our study, the calculation of the fitness $\lambda_{ind}$, was done following methods from McLean *et al.* (2019) and took into account all offspring born, regardless of their subsequent survival (whereas in the rest of our study, we only took into account offspring reaching sexual maturity). Because yellow baboons reproduce year-round in the wild (Campos *et al.*, 2017), we ran this internal validation with females following a non-seasonal phenology strategy.

The values simulated for major key life history traits (mass, fitness, mortality and interbirth interval) were very close to the values observed in the wild (Table S5), which tends to confirm our model's validity. However, the simulated values of body fat and post-partum amenorrhea (PPA) length were not as variable as in the natural population. This could partly be explained by the absence of a social hierarchy in the model, which is known to generate variation in many life history traits. Shorter maternal investment periods have indeed been reported for high-ranking females in several baboon species (Johnson, 2003; Alberts, 2019; Schneider-Crease *et al.*, 2022) and body mass was correlated with dominance rank in female chimpanzees (Pusey *et al.*, 2005). It is also possible that our model does not account precisely enough for the plasticity in offspring growth.

**2. Comparison of simulated and observed birth seasonality**

We evaluated, under realistic conditions, all 133 phenology strategies and subsequently selected the optimal ones (i.e. associated with the highest fitness values, Fig 3A, main text). These optimal phenology strategies were only slightly seasonal – as is typically the case of yellow baboons in the wild – with a minimal reproductive time window of 183 days (6 months) and included the non-seasonal phenology strategy.

We then computed the birth seasonality associated with the optimal phenology strategies by calculating the length of the mean vector resulting from all the births from these strategies and compared it to the value observed in the wild (Campos *et al.*, 2017). Simulated and observed birth distributions are presented in Figure S5. We found that simulated births resulting from pooling all optimal phenology strategies had a seasonality comparable ($r$ = 0.17) to the one observed in the wild ($r$ = 0.15). The mean birth date observed in the wild (10th Nov) occurred 3.5 months before the one computed from pooled optimal strategies (25th Feb). This slight increase in reproductive seasonality and delay in mean birth date could be due to some structural limitations of the model. For example, NDVI may not reflect exactly the real variation of food abundance experienced by an omnivorous species such as yellow baboons (Altmann, 1998) while the absence of social effects may also contribute to this discrepancy, as reproductive competition can lead to reduced birth seasonality in wild chacma baboons (Dezeure *et al.*, in press).

### TABLES

***Table S1: Description and values of the main parameters used in the model***

See Appendix B for justification and details about how parameters are used in the model.

| **Parameter** | **Description** | **Numerical value** | **Source or equation** | **Studied species** |
|---|---|---|---|---|
| Adult mass | Mean adult female total mass | 11.9 kg | Altmann *et al.*, 1993 | *Papio cynocephalus* (wild - Amboseli) |
| Age at beginning of weaning | Age at first significant consumption of plant food | 56 days | Rhine *et al.*, 1985 | *Papio cynocephalus* (wild - Mikumi) |
| Age at sexual maturity | Mean age at sexual maturity (first sex skin swellings) | 1643 days (4.5 years) | Charpentier *et al.*, 2008 | *Papio cynocephalus* (wild - Amboseli) |
| Birth mass | Mean birth mass | 0.710 kg | Altmann & Alberts, 1987 | *Papio cynocephalus* (wild - Amboseli) |
| Body fat increase duration | Average duration of the period of body fat increase before cycling | 30 days (1 month) | Gesquiere *et al.*, 2018 | *Papio cynocephalus* (wild - Amboseli) |

| | | | | |
|---|---|---|---|---|
| Cycling durations | Vector of observed cycling durations | Range: 18 - 590 days | Gesquiere *et al.*, 2018 | *Papio cynocephalus* (wild - Amboseli) |
| Energy in fat mass | Energy released by 1g of fat tissue | 39500 kJ/kg | Livesey & Elia, 1988 | *Homo sapiens* |
| Energy in lean mass | Energy released by or needed to create 1g of lean tissue | 20920 kJ/kg | Payne & Waterlow, 1971 | Mammals |
| Female lifespans | Vector of observed lifespans for breeding females | Range: 2234 - 9457 days | McLean *et al.*, 2019 | *Papio cynocephalus* (wild - Amboseli) |
| Gestation length | Mean gestation length | 178 days | Gesquiere *et al.*, 2018 | *Papio cynocephalus* (wild - Amboseli) |
| Growth rate | Mean weight gain by day during growth | 0.005 kg/day | Altmann & Alberts, 1987 | *Papio cynocephalus* (wild - Amboseli) |
| Infant proportion of external deaths | Proportion of infants dead from external causes | 0.1161 | Alberts, 2019; McLean *et al.*, 2019 | *Papio cynocephalus* (wild - Amboseli) |

| | | | | |
|---|---|---|---|---|
| Infant lifespans | Vector of observed lifespans for infants that died before weaning | Range: 1 - 319 days | McLean *et al.*, 2019 | *Papio cynocephalus* (wild - Amboseli) |
| Juvenile lifespans | Vector of observed lifespans for juveniles that died between weaning and sexual maturity | Range: 329 - 1731 days | McLean *et al.*, 2019 | *Papio cynocephalus* (wild - Amboseli) |
| Lactation efficiency | Efficiency of milk synthesis | 0.8 | Dewey, 1997 | *Homo sapiens* |
| Lean proportion | Mean lean mass proportion (of total mass) | 0.981 | Altmann *et al.*, 1993 | *Papio cynocephalus* (wild - Amboseli) |
| Longevity | Age of the oldest female reported at Amboseli | 9862 days (27 years) | Bronikowski *et al.*, 2002 | *Papio cynocephalus* (wild - Amboseli) |
| Mass at beginning of weaning | Estimated mass at beginning of weaning | 0.99 kg | = birthMass + growthRate× ageAtBeginningOfWeaning | |

| | | | | |
|---|---|---|---|---|
| Mass at Sexual Maturity | Approximate mean total mass at sexual maturity | 8.925 kg | = birthMass + growthRate× ageAtSexual Maturity | |
| Maximal daily fat storage | Maximal amount of fat that can be stored each day | 0.0001 kg | growthRate× (1-leanProportio n) | |
| Maximal energy intake | Maximal daily energy intake for a female baboon fed *ad libitum* | 4891.1 kJ/day | Roberts *et al.*, 1985 | *Papio cynocephalus* and *Papio anubis* (captive) |
| Maximal reduction of lean mass | Ratio of minimum lean mass causing death and regular lean mass | 0.85 | Payne & Waterlow, 1971; Scalfi *et al.*, 2002 | Mammals and *Homo sapiens* |
| NDVI | Vector of NDVI (Normalized Difference Vegetation Index) values in Amboseli (2000-2021) | Range: 0.13 - 0.42 | Didan, 2015 | |
| Placental proportion | Placenta mass expressed as a proportion of foetus mass | 0.25 | Farley *et al.*, 2009 | *Papio spp.* (captive) |

| | | | | |
|---|---|---|---|---|
| PPA duration | Mean postpartum amenorrhea duration | 322 day | Gesquiere *et al.*, 2018 | *Papio cynocephalus* (wild - Amboseli) |
| Storage efficiency | Efficiency of energy storage as fat mass | 0.9 | Rothwell & Stock, 1982 | *Rattus norvegicus* (Zucker rat - captive) |
| Weaning mass | Estimated threshold mass at complete weaning | 2.32 kg | = birthMass + growthRate× ppaDuration | |

***Table S2: Description of the main attributes for each class***

| **Attribute** | **Description** | **Owner(s)** | | | |
|---|---|---|---|---|---|
| *Mass* | | Foetus | Infant | Juvenile | Female |
| Total Mass (kg) | Total mass of the individual | ✔ | ✔ | ✔ | ✔ |
| Lean Mass (kg) | Lean mass of the individual | | ✔ | ✔ | ✔ |
| Fat Mass (kg) | Fat mass of the individual | | ✔ | ✔ | ✔ |
| *Condition* | | Foetus | Infant | Juvenile | Female |
| Alive | False if the individual is dead, true otherwise | ✔ | ✔ | ✔ | ✔ |
| Growth Allocation | 1 if growth is normal, 0 if growth is stopped and between 0 and 1 if growth is slowed. | | ✔ | ✔ | ✔ |
| *Timings* | | Foetus | Infant | Juvenile | Female |

| | | | | | |
|---|---|---|---|---|---|
| Cycle Time (d) | Day when the first cycle leading to the individual's conception begins | ✔ | ✔ | ✔ | ✔ |
| Conception Time (d) | Day when the individual is conceived | ✔ | ✔ | ✔ | ✔ |
| Birth Time (d) | Day of birth of the individual | | ✔ | ✔ | ✔ |
| Weaning Time (d) | Day when the individual is fully weaned | | | ✔ | ✔ |
| Sexual Maturity Time (d) | Day when the individual reaches sexual maturity | | | ✔ | ✔ |
| Death Time (d) | Day of death of the individual | ✔ | ✔ | ✔ | ✔ |
| Lifespan (d) | Life (or gestation) duration of an individual (or a foetus) before potential death (or miscarriage) from external causes | ✔ | ✔ | ✔ | ✔ |
| *Energy* | | Foetus | Infant | Juvenile | Female |
| Energy needs (kJ) | Energy needed to ensure maintenance, growth and potential reproduction of the individual | | | ✔ | ✔ |
| Energy intake (kJ) | Energy extracted from the environment | | | ✔ | ✔ |
| Energy released (kJ) | Energy released from storage (fat mass or lean mass) | | | ✔ | ✔ |
| Energy balance (kJ) | Difference between available energy (energy intake and energy released) and energy needs | | | ✔ | ✔ |
| *Reproduction* | | Foetus | Infant | Juvenile | Female |

| | | | |
|---|---|---|---|
| Phenology strategy (beginning and length of reproductive window) (d) | Time window during which reproduction is possible | | ✔ |
| List of Foetuses | Record of all ‘class Foetus’ individuals the female has produced | | ✔ |
| List of Infants | Record of all ‘class Infant’ individuals the female has produced | | ✔ |
| List of Juveniles | Record of all ‘class Juvenile’ individuals the female has produced | | ✔ |
| Cycling duration (d) | Duration of the ongoing cycling period | | ✔ |
| Reproductive status | Reproductive status of the female: "anoestrus", "oestrus", "gestation" or "lactation" | | ✔ |

***Table S3: Initial attributes for the female at the beginning of simulation (t=0)***

Unknown timings are given the value -999. Mass at sexual maturity, lean proportion and age at sexual maturity are given in Table S1.

| **Attribute** | **Initial value**<br>**Comments** |
|---|---|
| *Mass* | |
| Total Mass (kg) | massAtSexualMaturity = 8.925 kg |

| | |
|---|---|
| Lean Mass (kg)<br><br>Fat Mass (kg) | massAtSexualMaturity×leanProportion = 8. 755 kg<br><br>massAtSexualMaturity×(1-leanProportion) = 0.170 kg |
| *Condition* | |
| Alive<br><br>Growth Allocation | true<br><br>1 |
| *Timings* | |
| Cycle Time (d)<br><br>Conception Time (d)<br><br>Birth Time (d)<br><br>Weaning Time (d)<br><br>Sexual Maturity Time (d)<br><br>Death Time (d)<br><br>Lifespan (d) | -999<br><br>-999<br><br>-1643 = - ageAtSexualMaturity<br><br>-999<br><br>0<br><br>-999<br><br>Randomly picked from McLean *et al.*, 2019 |
| *Energy* | |
| Energy needs (kJ)<br><br>Energy intake (kJ)<br><br>Energy released (kJ)<br><br>Energy balance (kJ) | 0<br><br>0<br><br>0<br><br>0 |
| *Reproduction* | |
| Phenology strategy (beginning and length of reproductive window) (d)<br><br>List of Foetuses<br><br>List of Infants<br><br>List of Juveniles | Picked from the 133 different phenology strategies considered (see main text)<br><br>empty list<br><br>empty list<br><br>empty list |

| | |
|---|---|
| Cycling duration (d) | 0 |
| Reproductive status | anoestrus |

***Table S4: Initial attributes for offspring (foetus, infant and juvenile)***

The values given correspond to transitional events (conception, birth and weaning), where new individuals from associated classes (foetus, infant and juvenile) are created. A foetus is created from scratch at conception ($t=t_c$), but each infant is created from a foetus that has reached the end of gestation ($t=t_b$) and each juvenile is created from an infant that has reached the total weaning mass threshold ($t=t_w$). In the case where an individual is created from another one, it can take (inherit) some of its attribute values. Lean proportion is given in Table 2. See details for the "lifespan" attribute in the "death" submodel.

| **Attribute** | **Initial value** | | |
|---|---|---|---|
| | **Foetus (at conception: $t=t_c$)** | **Infant (at birth: $t=t_b$)** | **Juvenile (at weaning: $t=t_w$)** |
| | | *Mass* | |
| Total Mass (kg) | 0 | Inherited from foetus | Inherited from infant |
| Lean Mass (kg) | | totalMass x leanProportion | Inherited from infant |
| Fat Mass (kg) | | totalMass x (1-leanProportion) | Inherited from infant |
| | | *State* | |
| Alive | true | Inherited from foetus | Inherited from infant |
| Growth Allocation | | 1 | Inherited from infant |

| | | | |
|---|---|---|---|
| *Timings* | | | |
| Cycle Time (d) | Female's last beginning of cycle | Inherited from foetus | Inherited from infant |
| Conception Time (d) | $t_c$ | Inherited from foetus | Inherited from infant |
| Birth Time (d) | | $t_b$ | Inherited from infant |
| Weaning Time (d) | | | $t_w$ |
| Menarche Time (d) | | | -999 |
| Death Time (d) | -999 | Inherited from foetus | Inherited from infant |
| Lifespan (d) | 179 (gestationLength +1) or randomly picked between 1 and gestation length | Longevity or randomly picked from (McLean *et al.*, 2019) | Longevity or randomly picked from (McLean *et al.*, 2019) |
| *Energy* | | | |
| Energy needs (kJ) | | | 0 |
| Energy intake (kJ) | | | 0 |
| Energy released (kJ) | | | 0 |
| Energy balance (kJ) | | | 0 |

***Table S5: Comparison between simulated and observed data for key life history traits***

Mean value and standard deviation (mean ± sd) for simulated life history traits were obtained from 2000 simulations following the non-seasonal phenology strategy (reproduction was not constrained in time). The simulations were run using the NDVI extracted from Amboseli geological coordinates.

| **Life history trait** | **Observed data (mean ± sd)** | **Simulated data (mean ± sd)** | **Source** |
|---|---|---|---|
| Fitness ($\lambda_{ind}$) | 1.09 ± 0.08 | 1.09 ± 0.09 | McLean *et al.*, 2019 |
| Lifetime reproductive success | 5.61 ± 3.21 infants | 5.76 ± 3.16 infants | McLean *et al.*, 2019 |
| Age at death | 14.75 ± 5.42 years | 14.72 ± 5.42 years | McLean *et al.*, 2019 |
| Foetus Loss | 13.9% | 13.2% | Beehner *et al.*, 2006 |
| Infant Loss (before weaning) | 21.5% | 22.0% | Calculated from McLean *et al.*, 2019 |
| Offspring Loss (between birth and sexual maturity) | 45.3% | 45.0% | Calculated from McLean *et al.*, 2019 |
| Mean total mass | 11.9 ± 1.41 kg | 11.57 ± 0.32 kg | Altmann *et al.*, 1993 |
| Mean fat percent | 1.9 ± 4.81 | 2.2 ± 0.56 | Altmann *et al.*, 1993 |
| Interbirth interval duration | 638 ± 116 days | 637 ± 83 days | Gesquiere *et al.*, 2018 |
| Postpartum amenorrhea duration | 322 ± 87 days | 323 ± 17 days | Gesquiere *et al.*, 2018 |
| Cycling duration | 138 ± 82 days | 135 ± 80 days | Gesquiere *et al.*, 2018 |
| Gestation duration | 178 ± 6 days | 178 ± 0 days | Gesquiere *et al.*, 2018 |

### FIGURES

***Figure S1: Variation of energy intake with NDVI***

$E_{max}$ represents the maximum amount of energy an individual can get daily when food is non-limiting. For low values of NDVI, the energy intake associated increases rapidly, simulating the effect of “fallback food”. $\tau$ is the characteristic value of the equation and represents the speed at which $E_{max}$ is reached. The values of minimal, mean and maximal NDVI are given for the Amboseli national Park, and their associated energy intake are given as proportion of $E_{max}$.

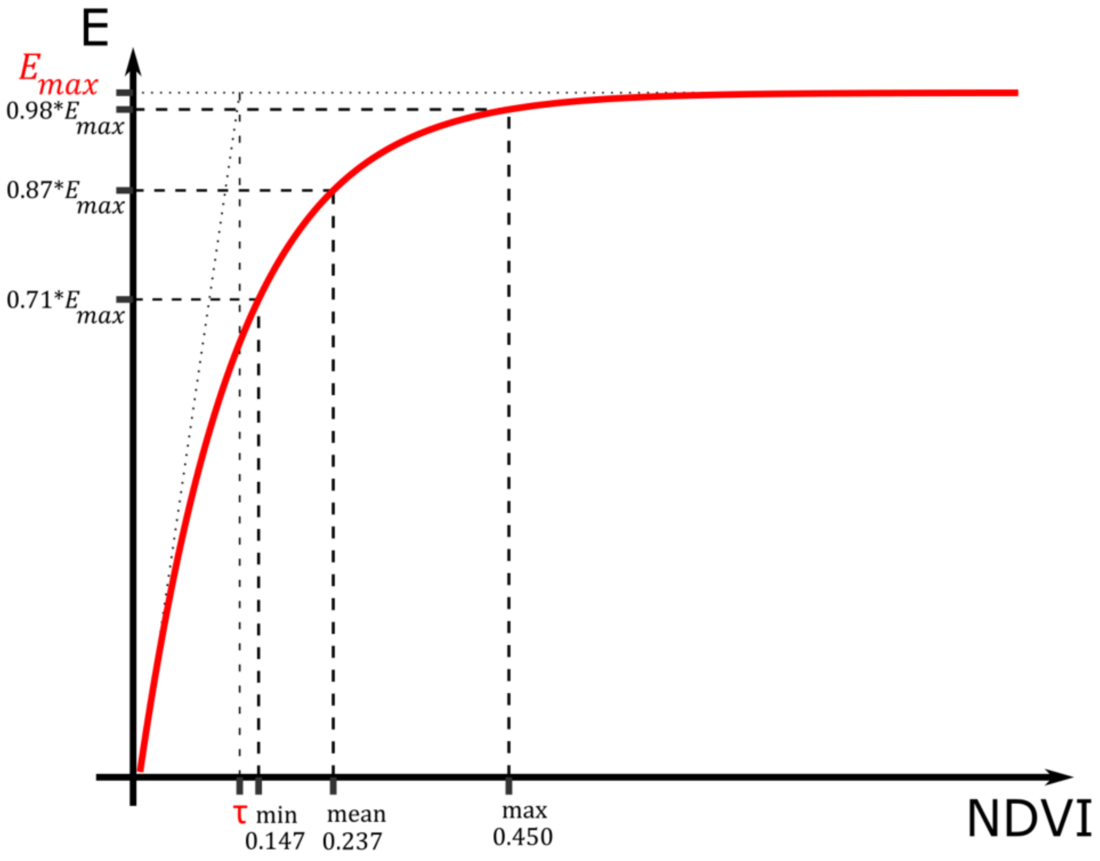

***Figure S2: Multiplying coefficient for energy intake during lactation (C)***

During lactation, energy intake is increased by a factor of 1.11 at birth, 1.27 at beginning of weaning and goes back to normal at the end of weaning. Weaning beginning and end are defined by threshold masses for the infant (Table 2).

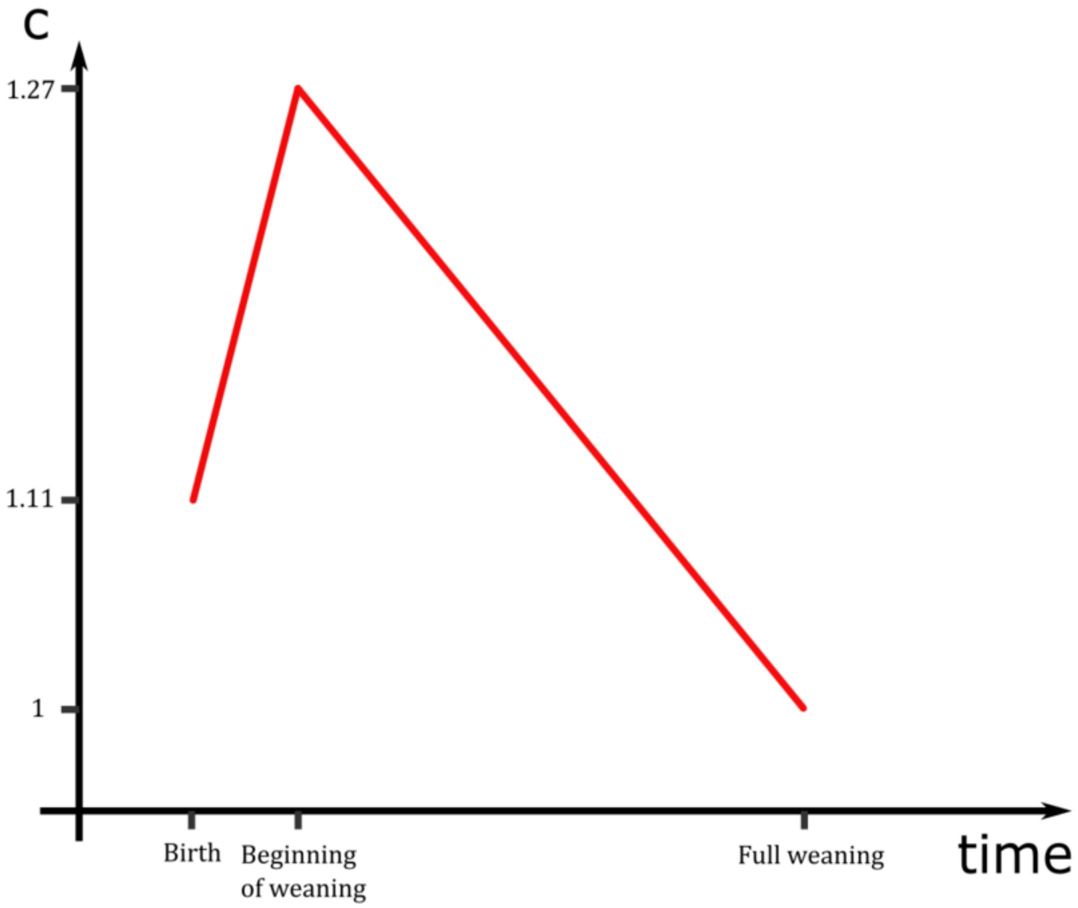

***Figure S3: Flowchart of energy allocation***

Energy balance (EB) is evaluated throughout the energy allocation decision tree.

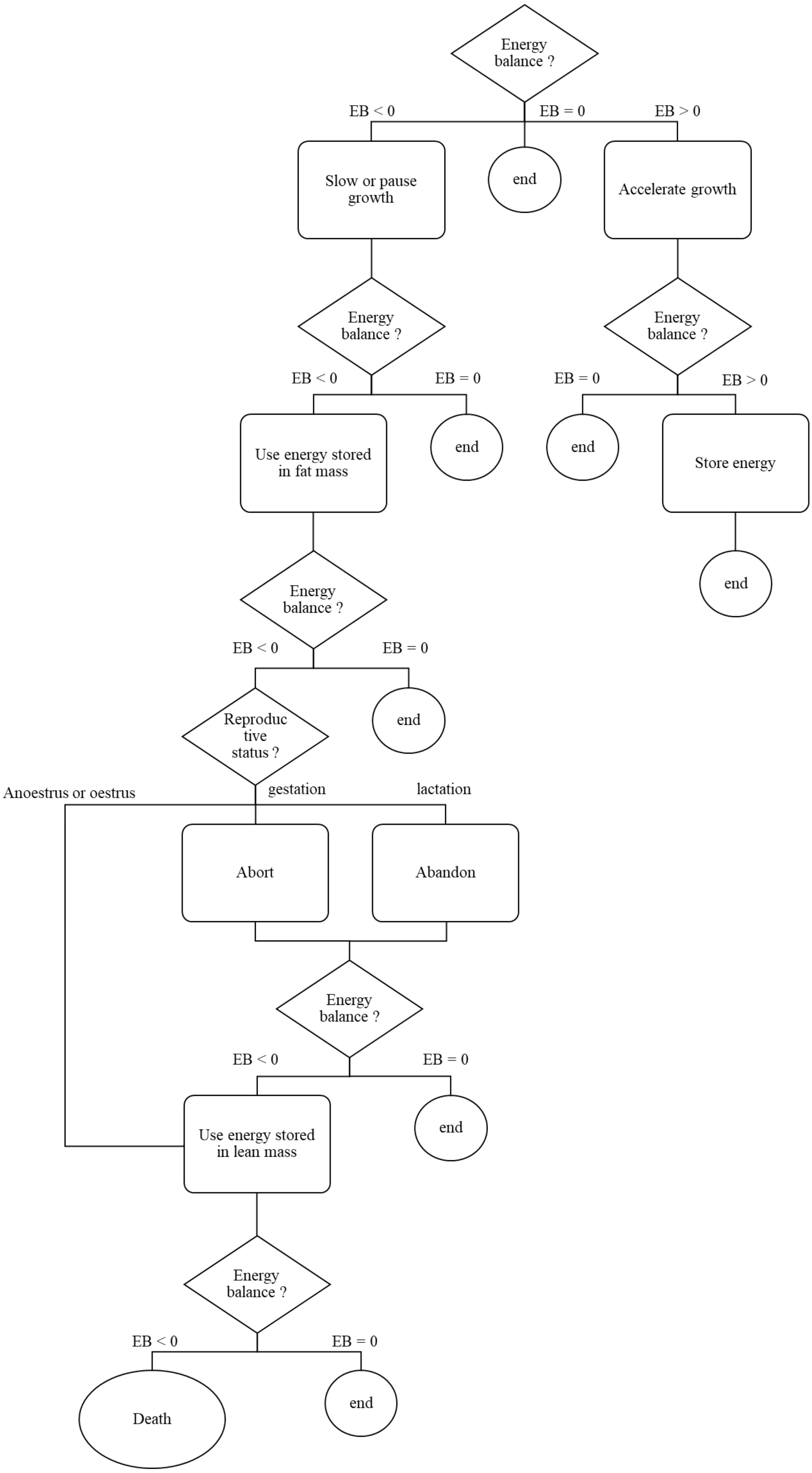

***Figure S4: NDVI time series decomposition***

The NDVI values were extracted according to the Amboseli geographical coordinates (South/West: -2.75, 37.04; North/East: -2.70, 37.11), and we only show ten years of data, from January 1st, 2001 to December 31st, 2010 for clarity purposes. Raw NDVI is plotted in the top panel, and is equal to the sum of the values plotted in the other three panels: the mean NDVI (K), the seasonal NDVI (NDVI_S) and the non-seasonal NDVI (NDVI_NS).

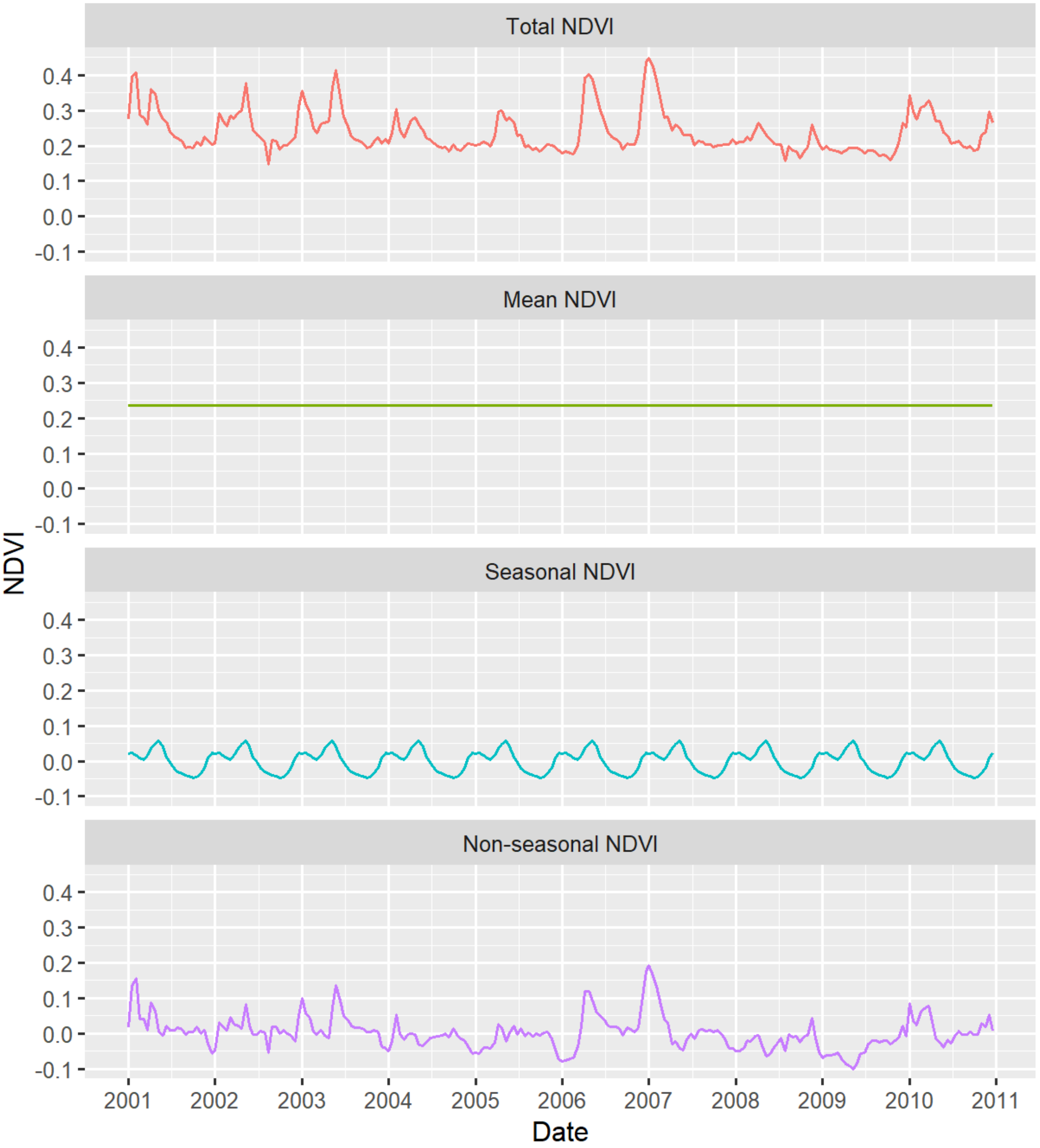

***Figure S5: Comparison of birth phenology***

Each circular box plot represents the monthly distribution of births and the red segment is the mean vector of births. Its direction, $\mu$, gives the mean date of birth while its length, $r$, quantifies the seasonality of births (r=0 means that births are equally distributed and r=1 means that all births occur on the same day of the year). The left panel shows the distribution of simulated births resulting from all the optimal phenology strategies (whose mean fitness were not significantly different). The right panel shows the real births distribution observed in the wild in Amboseli (Campos *et al.*, 2017).

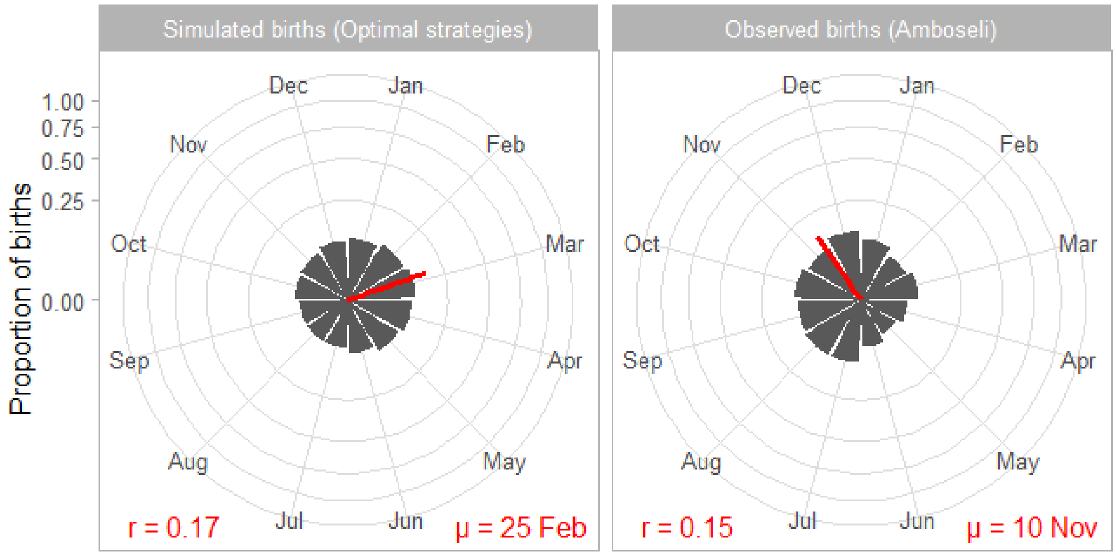

***Figure S6: Variation of mean individual fitness***

We plotted heatmaps of mean individual fitness ($\lambda_{ind}$) along gradients of environmental productivity and seasonality and with the different values we tested in this study for the four other factors: environmental unpredictability (u), daily reproductive energy expenditure (growth rate GR), reproductive cycle length (interbirth interval IBI as a function of year length YL) and infant extrinsic mortality rate (M). Associated birth seasonality (r = mean ± sd) is given below each heatmap. The top left panel represents the real conditions for these four factors, as observed in Amboseli.

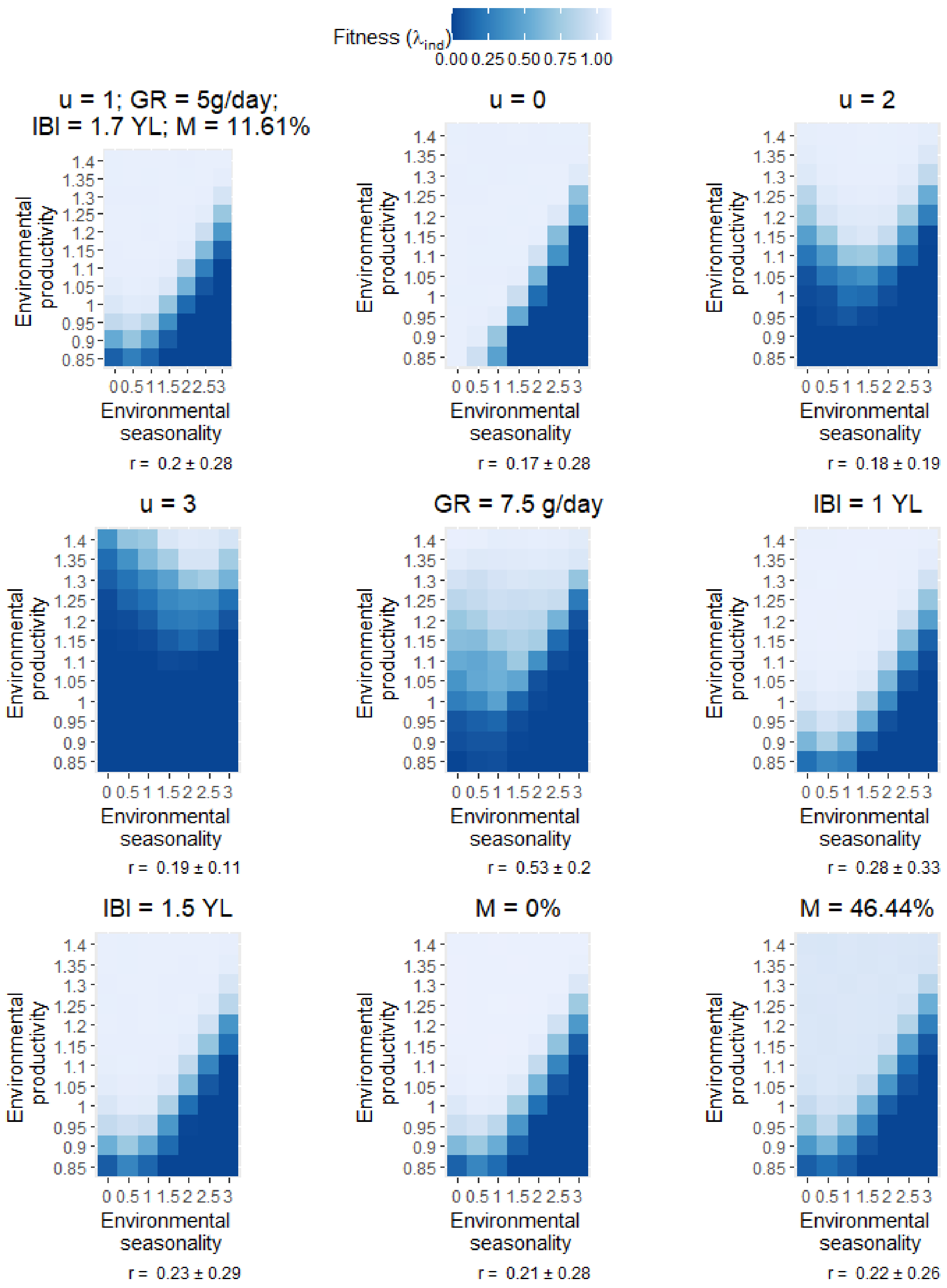
Fitness ($\lambda_{ind}$)
0.00 0.25 0.50 0.75 1.00
u = 1; GR = 5g/day;
IBI = 1.7 YL; M = 11.61%
u = 0
u = 2
u = 3
GR = 7.5 g/day
IBI = 1 YL
IBI = 1.5 YL
M = 0%
M = 46.44%
Environmental productivity
Environmental seasonality
r = 0.2 ± 0.28
r = 0.17 ± 0.28
r = 0.18 ± 0.19
r = 0.19 ± 0.11
r = 0.53 ± 0.2
r = 0.28 ± 0.33
r = 0.23 ± 0.29
r = 0.21 ± 0.28
r = 0.22 ± 0.26

***Figure S7: Effect of reproductive cycle length and infant mortality on birth seasonality (H5 and H6)***

Panel A shows the heatmaps of birth seasonality along gradients of environmental seasonality and productivity in three different conditions of reproductive cycle length: when the interbirth interval (IBI) is exactly one year long (1 YL), when its length is 1.5 years (1.5 YL), and in normal conditions, when it is 1.7 years long (1.7 YL). Panel B shows heatmaps of birth seasonality along gradients of environmental seasonality and productivity in three different conditions of infant extrinsic mortality: with no infant extrinsic mortality, with a regular infant extrinsic mortality of 11.61% and with a four-time amplified mortality of 46.44%. Birth seasonality is represented by r, going from 0 (births are equally distributed) to 1 (all births occur on the same day). Grey cells represent non-viable environments characterised by a fitness of zero (regardless of the phenology strategy followed), where r cannot be computed and generates missing values.

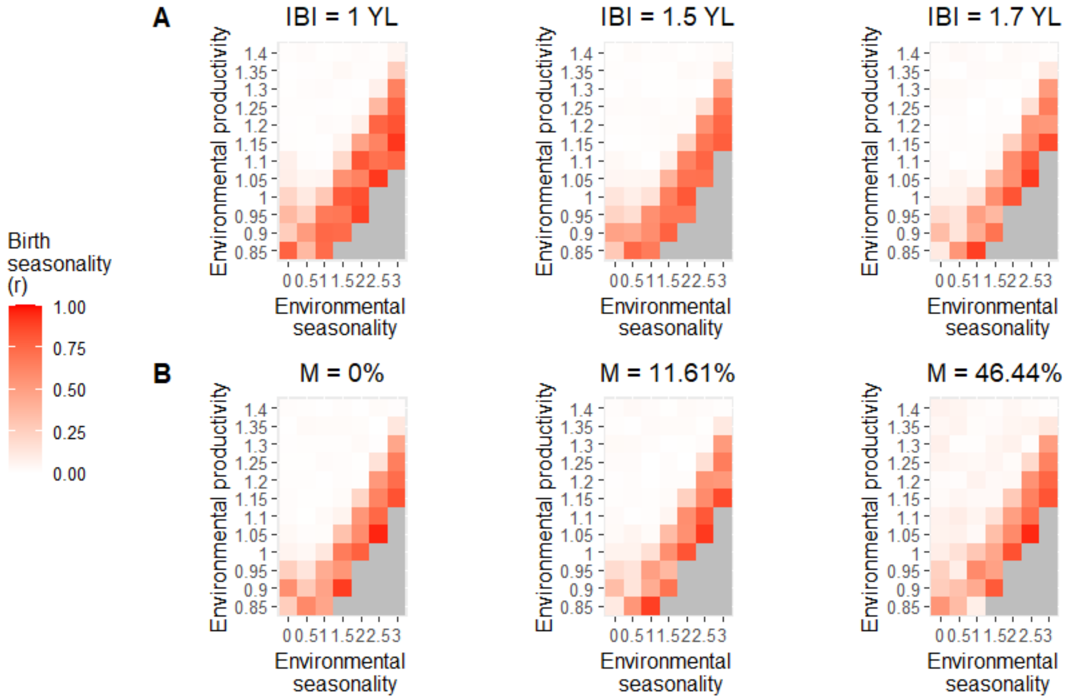
